# A convergent cortical scheme for visual scene segmentation

**DOI:** 10.64898/2025.12.28.696428

**Authors:** Minggui Chen, Huaixue Chen, Pieter R. Roelfsema, Wu Li

## Abstract

Scene segmentation, a key step towards structured understanding of natural environments, entails a complementary set of computations. Instead of accessing these segmentation computations in isolation, we perceive the segregated object as a unified whole, but the neural basis supporting such a unitary percept remains unclear. We bridged this gap by showing a convergent coding scheme: an object-level additive code in the primary visual cortex, which serves as a spatial segmentation mask for the object. Across individual neurons, this code uniformly boosted their responses to the segmented object. In the population space, this code created an orthogonal representational geometry by perpendicularly displacing the neural manifold for image features, efficiently multiplexing both local and global information about the object. Computational modeling further revealed the utility and optimality of this convergent coding scheme, which effectively isolates objects in cluttered environments and in turn unclutters downstream object representations. Our findings reveal the computational logic of visual segmentation and call for a revision of the visual cortical hierarchy.

---

Our visual system starts parsing natural scenes by discretizing retinal images into local visual features (*1*). The fragmented information extracted by such low-level processing needs to be organized in a structured manner for proper interpretation of the visual scenes (*2–4*). An effective solution offered by a series of mid-level processes, collectively known as perceptual grouping and segmentation, imposes global structures onto the local features (Fig. 1A, two middle panels), integrating elements belonging to the same object and segregating them from the background elements (*5, 6*). These processes are crucial for high-level visual behaviors including object recognition (*7*).

**Fig. 1.**
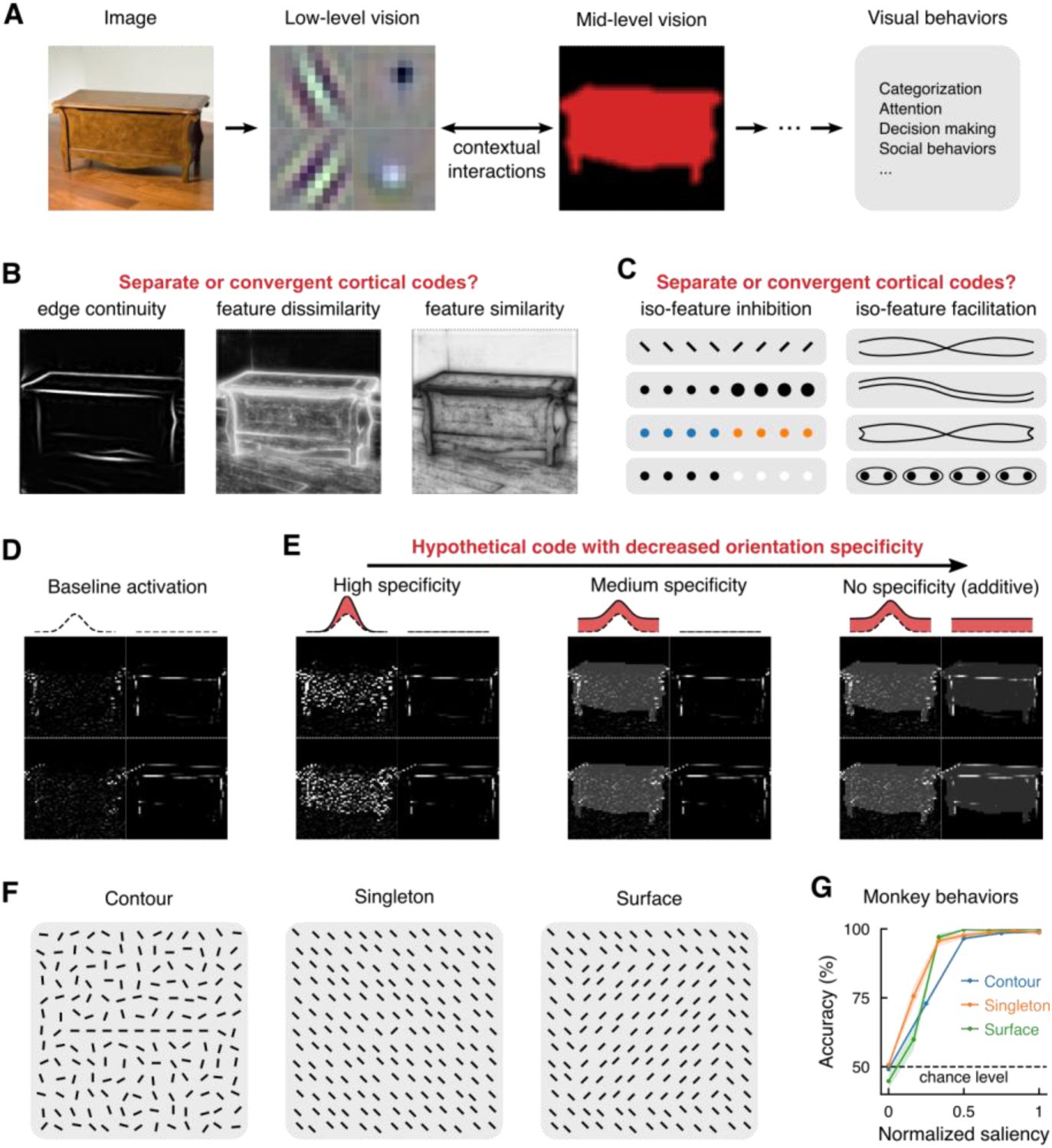
Hypotheses and paradigms. **(A)** Hierarchical stages of image analysis. An image is analyzed by local orientation-selective and orientation-nonselective filters during the low-level visual process. In the mid-level process, a complementary set of computations imposes structured contextual information onto the low-level process to group and segment visual features, which in turn supports high-level visual processes and visually-guided behaviors. **(B)** Effects of three complementary segmentation computations (see methods). The computation for finding edge continuity boosts the signal-to-noise ratio of continuous contours; the computation for detecting feature dissimilarity defines the boundaries between visual features; and the computation for detecting feature similarity helps to partition the image into separate regions. How these computations collectively give rise to a unitary segmented object (e.g., a desk in the room) remains largely unknown. **(C)** Gestalt principles of perceptual organization sorted into two categories by two opposite computations: Iso-feature inhibition detects abrupt changes in visual features whereas iso-feature facilitation integrates continuous contour segments and assembles similar image components, leading to perceptual grouping and segmentation of image components. **(D)** Baseline activations of 4 individual visual filters. The sample image in A was convolved, respectively, with the two orthogonal orientation-selective filters shown in A (second panel, left column) and the two orientation-nonselective filters (right column) to visualize their outputs. The insets on top schematize the orientation tuning curves of these two types of filters. **(E)** Visualization of three hypothetical segmentation codes with different degrees of specificity (illustrated as the red areas in top insets), which resulted in different modifications to the baseline activations shown in D (see text and methods for details). **(F)** Stimulus designs for the segmentation experiments. Contours, singletons, and surfaces were respectively used to study the three complementary computations shown in panel B (i.e., edge continuity, feature dissimilarity, and feature similarity). **(G)** The animal’s behavioral performance on detection of the 3 types of simple objects shown in F at different levels of object saliency. Results were pooled from the two monkeys and the saliencies of different objects were rescaled within 0-1. See fig. S1 for sample stimuli of different saliencies and corresponding behavioral performance.

Behavioral and theoretical investigations (*8*), as early as the Gestalt psychology (*5, 6*), have identified a complementary set of principles and computations underlying the segmentation of visual objects (*9–11*). For instance, object boundaries are usually delineated by integrating smoothly arranged contour segments (Fig. 1B, left) and by detecting abrupt changes of visual features (Fig. 1B, middle). After demarcation of the object boundaries, the interior region of the object is filled in (Fig. 1B, right). A combination of these computations not only explains the classical Gestalt laws derived from simplified artificial stimuli (Fig. 1C), but also accounts for various visual effects in segmenting natural scene images (*5, 12, 13*). In line with the visual effects, these segmentation computations are initiated in different cortical circuits, as evidenced by the differences in the progression of neural activities in the striate and extrastriate cortices (*14–16*) and by causal manipulations of the extrastriate cortex (*17–20*). These specialized neural processes eventually give rise to a unitary percept of the segmented object (*4, 21*), but it remains unknown whether these neural processes for image parsing would converge onto a unified cortical scheme for image segmentation.

At the level of individual visual filters (e.g., Fig. 1A, 2nd panel), image segmentation could be implemented by differentially modulating the original baseline activations of assorted filters (Fig. 1D; see methods for details). For example, by multiplicatively (Fig. 1E, left panel) or additively (Fig. 1E, middle) uplifting the response gain of orientation-selective filters within the object region, the oriented image components belonging to the object will be highlighted, with the modulation strength dependent on (Fig. 1E, left), or irrespective of (Fig. 1E, middle), the degree of orientation matching between the filters and the image components. Alternatively, the activities of all kinds of filters within the object region could be uniformly boosted regardless of the filters’ feature selectivities (Fig. 1E, right). Given these possible forms of the segmentation code with different degrees of specificities, what form would be taken by a convergent coding scheme for image segmentation remains unknown.

To address the above questions, we utilized three representative stimulus designs for visual segmentation in the orientation domain (Fig. 1F). The first stimulus was an elongated contour standing out from the background because of collinearly aligned line elements. The second stimulus contained a singleton: a bar popping out from the background due to its unique orientation. The third stimulus was a square surface segregated from the background as a result of a difference in texture orientations. By probing into the neural computations for detecting edge continuity, feature dissimilarity, and feature similarity (see Fig. 1B), we sought to investigate: (i) whether there is a convergent neural code for image segmentation, (ii) the format of the code, (iii) how the codes for local features and for image segmentation coexist at the neural population level, and (iv) the computational logic of segmentation.

## Behavioral paradigms and neuronal recordings

Unless stated otherwise, the visual display consisted of short bars (0.25° × 0.05°) distributed in an invisible grid of 0.5° squares for the three segmentation scenarios examined (Fig. 1F and fig. S1; see methods). Two macaque monkeys were trained to segment a simple object (contour, singleton, or surface) from the background. For the contour object (Fig. 1F, left), its saliency was determined by the number of collinear bars (1 to 9 bars) in a background of randomly oriented bars (fig. S1A). For the singleton (Fig. 1F, middle) and surface (9 × 9 parallel bars; Fig. 1F, right), the saliency of the foreground object depended on the angular difference (0-90°; termed the orientation contrast) between the object and background bars (fig. S1, B and C). In a trial (fig. S2), the stimulus display lasted 500 ms and the object was randomly embedded in the background at one of two predefined locations symmetrical about a fixation spot. After the stimulus disappeared, the monkey was required to make a saccade to indicate the object location. For the three types of stimuli, the animals’ behavioral performance significantly increased with object saliency (Fig. 1G; Pearson correlation: all *ρ*s > 0.93, all *P*s < 10^−3^).

While the monkey was performing the segmentation tasks, we recorded neuronal spiking activities in the primary visual cortex (area V1) with chronically implanted microelectrode arrays (see methods). In a trial, the receptive field (RF) of the neurons recorded by an electrode (referred to as a V1 site) was centered either on the central component bar of the object (termed the object-centered condition; fig. S3, 1st column) or on a bar in the background (the object-absent condition; fig. S3, 2nd column). The responses of a V1 site (100-550 ms after stimulus onset) were averaged over the trials in the same stimulus condition and then normalized across different conditions. The difference of normalized mean responses between the object-centered and object-absent conditions was defined as the segmentation signal. The segmentation signals were analyzed for the most salient objects (contour formed by 9 collinear bars; singleton or surface with 90° orientation contrast). The V1 sites from the two animals in the same experiment were pooled as the results were qualitatively similar (see scatter plots where individual sites from the two animals are indicated by light and dark colors, respectively).

The recorded V1 sites were classified into orientation selective or nonselective groups based on their responses to grating stimuli of different orientations (see methods). The optimal orientation of a V1 site was determined from the orientation tuning peak. The degree of orientation selectivity of a V1 site was measured as the ratio of the mean neuronal responses to orthogonally and optimally oriented gratings after subtracting the mean spontaneous activity.

## A convergent segmentation code in V1

Although V1 recording sites showed a broad range of selectivity for grating or bar orientations, from highly selective to nonselective, the segmentation signals for the global contour (Fig. 1F, left) were independent of the selectivity. Take the contour segmentation as an example. When the orientation tuning curves of individual V1 sites in the object-centered condition were contrasted with those in the object-absent condition (fig. S3, A to E; in both conditions one bar was always centered in the RF, see fig. S3A), the contour segmentation signals were comparable across stimulus orientations no matter whether a V1 site was orientation selective or nonselective. The mean segmentation signals averaged across stimulus orientations were also comparable between the selective and nonselective neuronal populations (Fig. 2A). A lack of interaction between orientation tuning and the segmentation signal was further confirmed by the absence of a correlation (Fig. 2B; *ρ* = −0.01, *P* = 0.91). When we aligned and averaged the tuning curves for the orientation-selective and nonselective sites separately (see methods), we observed that the presence of the object on the RF increased the responses of both groups of sites in an additive manner and by a comparable amount (Fig. 2C). The bell-shaped tuning curves of orientation-selective sites were fitted by Gaussian functions with four parameters—the amplitude, the optimal orientation (the Gaussian center), the tuning bandwidth (the standard deviation), and the pedestal on which the Gaussian was superimposed. The collinear contour lying on the RF increased the pedestal and the amplitude (Fig. 2D; all *P*s < 10^−7^), leaving other parameters unchanged (Fig. 2E; all *P*s > 0.13). Further analyses showed that the pedestal elevation largely accounted for the overall change in the tuning curve (77%, Fig. 2F; see methods for details). These results indicate that the presence of the contour object added a relatively constant number of spikes to a neuron’s responses, producing a general additive code independent of the neuron’s orientation tuning.

**Fig. 2.**
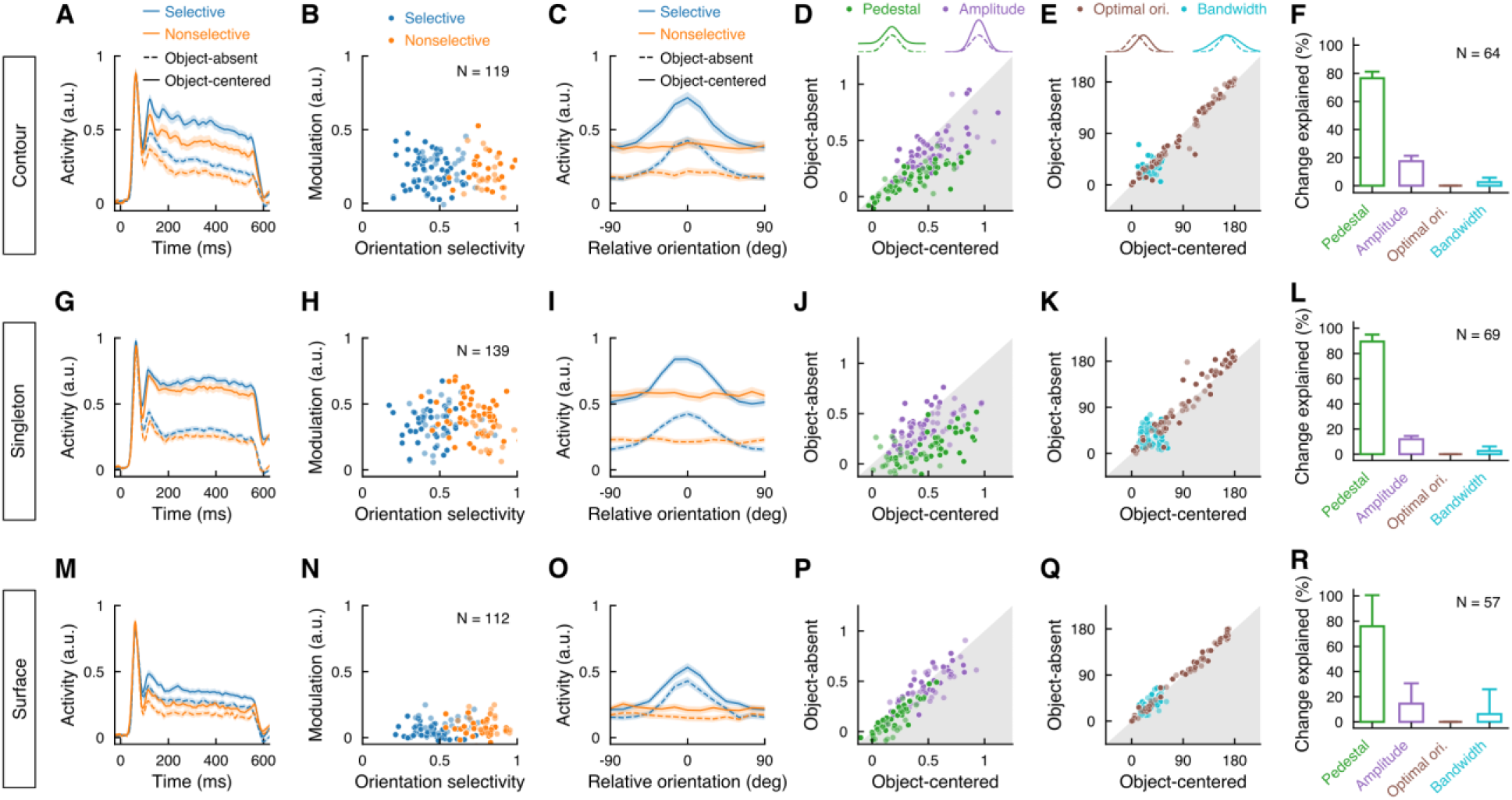
The convergent additive code in V1 for image segmentation. Three rows correspond to the three types of stimuli indicated on the left. **(A)** Population PSTHs (peri-stimulus time histograms; see methods) in response to the contour stimuli in the object-absent (dashed) and object-centered (solid) conditions, averaged across all the contour orientations and plotted separately for orientation-selective (blue) and nonselective (orange) V1 sites. Color shades indicate standard errors of the mean (SEM); time 0, stimulus onset. **(B)** Comparison between the strengths of segmentation signals (i.e., modulation) and orientation selectivities (see definitions in the text and methods) across individual V1 sites. Light and dark points represent data from the two animals, respectively (same in other scatter plots). **(C)** Population averaged orientation tuning curves in the object-absent and object-centered conditions for orientation-selective and nonselective V1 sites, respectively. The tuning curves of individual V1 sites were aligned to their respective optimal orientations (see methods). **(D** and **E)** Comparisons of Gaussian fitted orientation tuning parameters between the object-absent and object-centered conditions for orientation-selective sites. Each pair of Gaussians in top insets illustrate the tuning curves when the corresponding tuning parameter is increased in the object-center condition (solid) relative to the object-absent condition (dashed). **(F)** Respective contributions of individual tuning parameters to the overall changes in tuning curves between the object-absent and object-centered conditions (see methods). **(G** to **R)** Companion visualizations for the singleton (G to L) and surface (M to R) experiments. See fig. S4 for similar results for the streaked surface.

Striking similarities were observed among different segmentation scenarios (Fig. 1F), as shown in individual examples of V1 sites (fig. S3) and across the entire population (Fig. 2) in response to the contour, singleton, and surface. The similarities were manifested in comparable segmentation signals between the orientation-selective and nonselective neurons (Fig. 2, A, G and M), a lack of correlation between the segmentation signals and orientation tuning (Fig. 2, B, H and N; absolute values of all *ρ*s < 0.13, all *P*s *>* 0.15), and a vertical shift of the tuning curve (Fig. 2, C, I and O) mainly caused by an increase in the tuning pedestal (Fig. 2, D to F, J to L, and P to R; for the pedestal increment, all *P*s < 10^−7^). Across the orientation-selective sites, we also observed a small but significant increase in the tuning amplitude for the contour and singleton stimuli (Fig. 2, D and J, purple; all *P*s < 10^−5^) but not for the surface stimuli (Fig. 2P; *P* > 0.09). Nevertheless, for all the three segmentation scenarios, the apparent changes in orientation tuning curves of V1 neurons were largely explained by the additive increase (76%-90%; Fig. 2, F, L, R). A similar additive code was also observed for another popular design of surface segmentation (referred to as the streaked surface; see fig. S4).

The above results indicate a convergent coding scheme for image segmentation, which reconciles many previous observations of comparable temporal (*14, 22–27*) and spatial (*15, 28–30*) patterns of V1 responses to complex figure-ground stimuli. The additive segmentation code might provide an efficient and universal mechanism for image segmentation, highlighting all the visual features confined to the object region and rendering the entire object conspicuous on the cluttered background. This conjecture of object-level saliency was confirmed by the following experiments.

## The segmentation code and object-level saliency

We shifted the contour or surface formed by optimally oriented bars so that the central or a distal component bar of the object was centered in the RF of the same V1 site (Fig. 3, A and D). In accordance with the conjecture of object-level saliency, the enhanced neuronal responses were independent of the RF locations on the object (Fig. 3, B and C for the contour, *P* = 0.15; Fig. 3, E and F for the surface, *P* = 0.62).

**Fig. 3.**
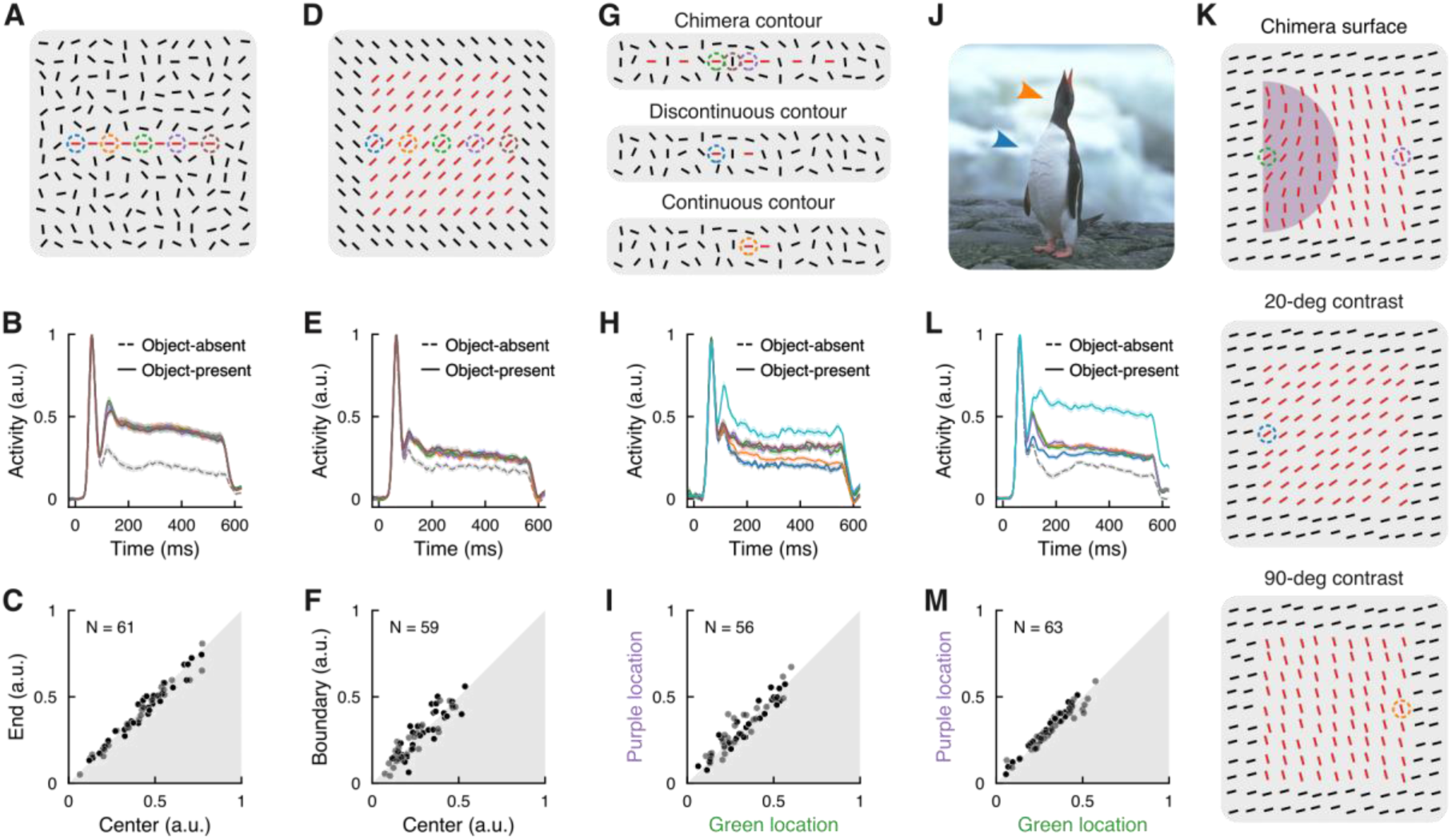
The segmentation code and object-level saliency. **(A)** Responses of the same orientation-selective V1 site were sampled respectively by placing the RF (circle) at 5 equally spaced locations along the optimally oriented contour. **(B)** Population PSTHs at the 5 locations on the contour (cluster of 5 solid curves) and in the object-absent condition (dashed). **(C)** Comparison of individual V1 sites’ responses to the component bar at the center and the end of the contour (two ends were pooled). **(D** to **F)** Companion visualizations for the surface stimuli. **(G)** Illustration of the chimera contour (top), which was formed by concatenating two short contour fragments (a triplet of two collinear bars with an orthogonal bar in between, middle; and a pair of adjacent collinear bars, bottom) and then appending additional bars aligned and non-aligned with the contour path (see methods). Neuronal responses were measured by placing the RF at the designated locations (circles in different colors) on the chimera contour or the contour fragments. Two complementary stimulus sets were pooled to balance the orientation of the component bar in the RF: The global contour path (and thus the component bar in the RF) was either aligned with, or orthogonal to, the optimal orientation of the recorded V1 site (fig. S5A. top row). **(H)** Population PSTHs in the object-absent condition (the lowest gray curve behind the blue one) and at the 5 locations shown in G (circle colors match the PSTHs). Note the superimposition of the PSTHs at the 3 locations on the chimera contour. The highest PSTH (cyan) represents a control condition in which all bars were collinearly aligned (fig. S5A, bottom row). **(I)** Comparison of individual V1 sites’ responses to the component bar at the purple and green locations on the chimera contour. **(J)** A natural image illustrating object contours with different local saliencies (e.g., arrow-pointed locations). **(K)** The chimera surface (top) was created as follows (see methods): The bars belonging to the surface but outside the semicircular area were set at the largest orientation contrast of 90° relative to the background (compare top and bottom panels); and the orientation contrast of the bars within the semicircular area gradually reduced, starting from 90° along the semi-circumference and decreasing to 20° at the diameter center (compare top and middle panels). Two complementary stimulus sets were pooled to balance the orientation of the component bar in the RF (fig. S5B, top and middle rows): The surface component bar in the RF was optimally or orthogonally oriented, keeping its orientation contrast unchanged relative to the background. **(L)** Population PSTHs in the object-absent condition (the lowest PSTH, gray) and at the 4 locations shown in K (circle colors match the 4 PSTHs). The singleton stimulus (fig. S5B, bottom row) was used as a control (the highest PSTH, cyan). **(M)** Comparison of individual V1 sites’ responses to the two boundary locations on the chimera surface, where the local orientation contrasts (relative to the background) differed by 70° (see K, top). The contours and surfaces in the sample stimuli are highlighted in red simply for illustrative purposes.

The location-invariant segmentation signal, or the object-level saliency effect, was seen even after we introduced heterogeneity in the component bars forming the foreground contour and surface.

We first created an elongated contour that was interrupted by non-aligned bars (Fig. 3G, top, termed the chimera contour; see methods). Two smaller fragments of the chimera contour, one consisting of two collinear bars interrupted in between by an orthogonal bar (Fig. 3G, middle) and the other consisting of two adjacent collinear bars (Fig. 3G, bottom), elicited significantly different responses due to different local saliencies (Fig. 3H, blue vs. orange curves; *P* < 10^−7^). However, when the two contour fragments were concatenated into the longer chimera contour (Fig. 3G, top), neuronal responses to them were raised to an identical level (Fig. 3H, overlapped green and purple curves; Fig. 3I, *P* = 0.74). The same level of response was reached even if the RF was centered on the component bar orthogonal to the contour path (Fig. 3G, top, middle bar in the brown circle; Fig. 3H, barely visible brown curve superimposed on the purple and green ones). The uniform response enhancement along the chimera contour was not caused by a ceiling effect, because aligning all elements on the contour path (fig. S5A, bottom row) further increased the neuronal responses (Fig. 3H, cyan). The chimera contour is reminiscent of contours in natural scenes (Fig. 3J), in which contour segments of an object with low and high local saliencies are perceived as an integrated whole.

Similar results were observed for segmentation of chimera surfaces with varying local orientation contrast (Fig. 3K, top; see methods). The RF of an orientation-selective V1 site was centered on opposite boundaries of the chimera surface with a low or high local orientation contrast relative to the adjacent background bars (20° vs 90°; compare the green and purple circles in Fig. 3K, top). As a comparison, we also presented two homogeneous surfaces having a constant orientation contrast of 20° and 90°, respectively (Fig. 3K, middle and bottom). For the two homogeneous surfaces, the one with the lower orientation contrast elicited the weaker response (Fig. 3L, blue vs. orange; *P* < 10^−8^). Interestingly, for the chimera surface, the responses of the same V1 site to the bars with the low and high local orientation contrasts (20° vs. 90°, Fig. 3K, top) became indistinguishable (Fig. 3L, overlapped green and purple curves; Fig. 3M, *P* = 0.13), indicating the uniformity of the surface segmentation signal.

All the observations above consistently point to an object-level segmentation code whereby all the locations occupied by the object are uniformly highlighted, resulting in a saliency effect at the object level.

## Object-level attributes other than saliency

Our visual system not only segments different objects but also analyzes their overall shapes (*31*). To examine whether V1 encodes object-level attributes such as global orientations, we set the collinear contour at either the optimal or orthogonal orientation of a V1 site while maintaining the local stimulus identical within the RF (Fig. 4A). Comparable neuronal activities and thus similar segmentation signals were observed for the two orthogonal object orientations (Fig. 4B for population averaged PSTHs; Fig. 4C for mean responses of individual sites, *P* = 0.72). The same neurons, however, showed a striking difference when the local bar in the RF took the optimal and orthogonal orientations (fig. S6, A to C). Similar results were obtained when testing with a slim version of the surface (Fig. 4D; see methods): V1 neurons were insensitive to changes in the global surface orientation (Fig. 4, E and F; *P* = 0.44), generating constant segmentation signals; but the same neurons showed typical bell-shaped tuning curves to the local bar orientations (fig. S6, D to F). These results suggest that V1 neurons play a dual role, labeling local visual features and signaling their object-level saliencies, irrespective of the global orientation of the object.

**Fig. 4.**
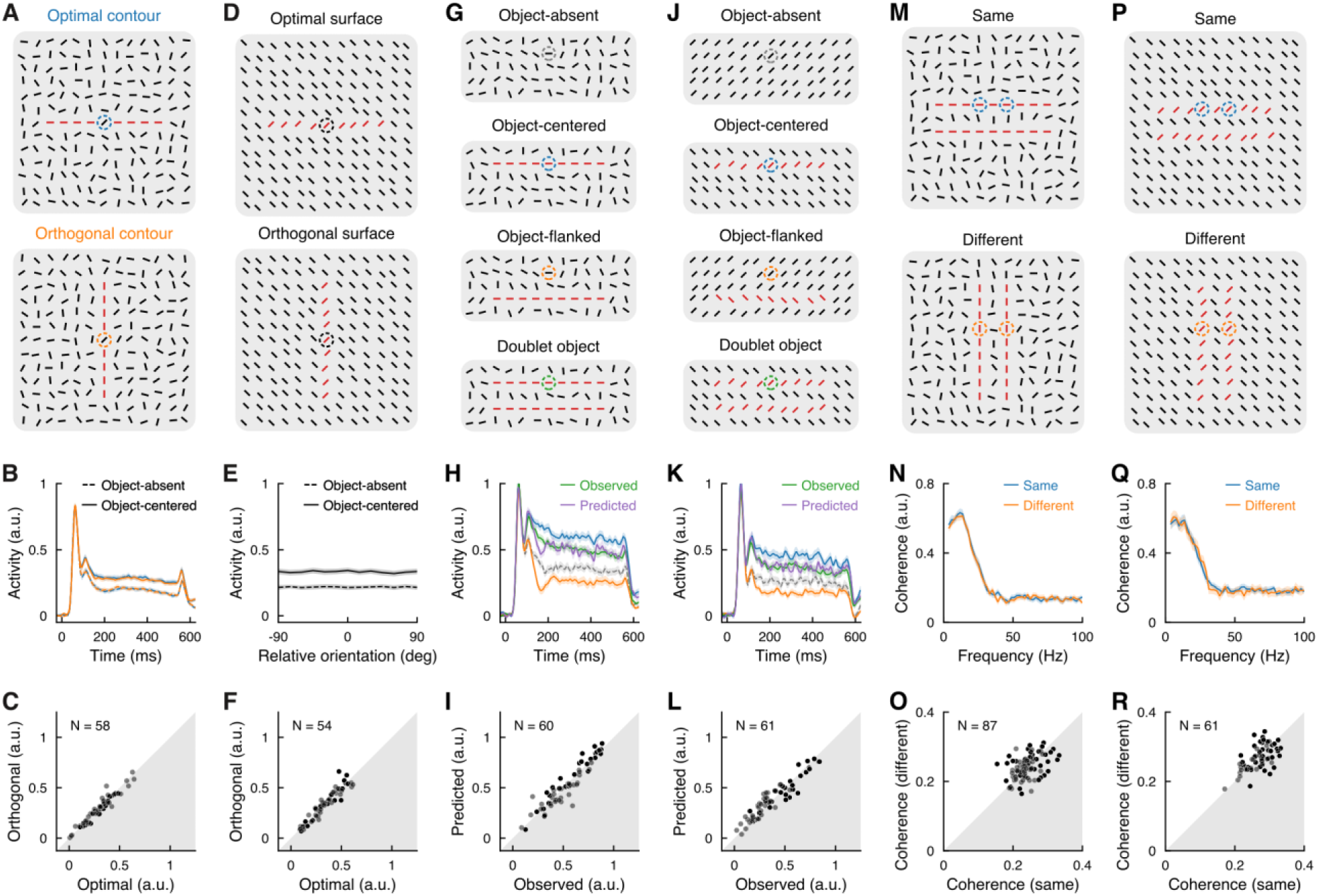
Object-level attributes other than saliency. Only orientation-selective sites were included. **(A)** The contour path was either aligned with, or orthogonal to, the optimal orientation of the recorded V1 site, keeping the bar in the RF unchanged. **(B)** Population PSTHs at the optimal (blue) and orthogonal (orange) contour orientations in the object-absent (dashed) and object-centered (solid) conditions. **(C)** Comparison of individual V1 sites’ mean responses to the optimally and orthogonally oriented contours shown in A; same data as in B. **(D)** The global orientation of the slim surface was varied by rearranging the orientation of the hidden grid used for placing the bars (see methods). A total of 12 global orientations (0-180°, 15° apart; two examples are shown here) were tested together with the 12 corresponding conditions in the absence of the slim surface (i.e., object-absent conditions). For each global orientation, a total of 12 local orientations of the surface bars (0-180°, 15° apart; an example of 45° is shown here) were tested and then pooled. The orientation contrast between the surface and background bars was always kept at 90°. **(E)** Population-averaged tuning curve for the slim-surface orientation (solid), which was obtained by aligning the tuning curves of individual V1 sites to their respective optimal orientations (see methods). Corresponding curve in the object-absent condition is also shown (dashed). Despite a lack of selectivity for global object orientations, the same neurons were selective for local bar orientations (fig. S6E). **(F)** Comparison of individual V1 sites’ mean responses to the optimally and orthogonally oriented slim surfaces. **(G)** Contour stimuli for testing the linear prediction of neuronal responses to multiple objects. **(H)** Population PSTHs in the object-absent (gray), object-centered (blue), object-flanked (orange), and doublet object (green) conditions illustrated in G. Purple curve indicates the predicted PSTH in the doublet object condition based on a linear sum of response components induced by individual contours. **(I)** Comparison between the observed and predicted mean responses of individual V1 sites in the doublet object condition shown in G; same data as in H. **(J** to **L)** Same as G to I, for the slim-surface stimuli. **(M)** Contour stimuli for examining inter-neuronal interactions, illustrating a pair of V1 RFs (circles) on the same object (top) or different ones (bottom). **(N)** Spike-spike coherence averaged over all recorded pairs of V1 sites. **(O)** Comparison of the mean coherences (0-100 Hz as in N) of individual pairs of V1 sites between the two conditions shown in M. **(P** to **R)** Same as M to O, for the slim-surface stimuli.

Next, we asked, in the presence of multiple objects, whether the segmentation code in V1 still operates in a simple additive manner. We compared three stimuli embedded with 0, 1, and 2 collinear contours (see methods). Compared with the object-absent condition (Fig. 4G, top), a single contour placed on the RF (Fig. 4G, 2nd panel) or at a location 1° from the RF center (Fig. 4G, 3rd panel) enhanced or suppressed the neuronal responses (Fig. 4H, blue vs. gray and orange vs. gray; all *P*s < 10^−8^). In the presence of two contours, one on the RF and the other at the location 1° aside (Fig. 4G, bottom), the responses were well approximated by a linear sum of the above enhancement and suppression (Fig. 4H, green and purple; Fig. 4I; *P* = 0.35). The linear prediction also held in the presence of two slim surfaces (Fig. 4, J to L; predicted vs. observed responses, *P* = 0.20), suggesting independent coding of multiple objects.

To examine whether multiple object representations involve neural synchronization, we compared the coherence (see methods) between the spike trains of a pair of V1 sites when their RFs were located on the same object or different ones (Fig. 4, M and P). There was no difference between the same- and different-object conditions (Fig. 4, M to R; all *P*s > 0.16), indicating that the additive segmentation code in V1 operates at the level of individual objects.

## A convergent geometry of the segmentation manifold

The series of experiments in the current study showed the coexistence of two types of neural codes in V1: the selective code for visual features confined to the RF and the general code for visual segmentation at the object level. How are these two distinct codes reconciled in the same population of neurons? This is crucial for downstream areas to derive a veridical object representation from the mixed V1 outputs. One hypothesis would be multiplexing through orthogonal codes so that potential interferences between them could be minimized. The additive code seen in individual V1 sites, i.e., a similar modulation across a wide range of firing rates in response to different local orientations, implies an orthogonality at the single neuron level. We further tested this hypothesis at the population level.

Based on each dataset obtained with a particular stimulus type (contour, singleton, or surface in Fig. 2; streaked surface in fig. S4), we trained a linear classifier (Fisher linear discriminant, FLD; Fig. 5A; see methods) to discriminate between two adjacent orientations of the bar in the RF (the feature decoder) or between the object-centered and object-absent conditions (the object decoder). Both types of decoders were trained separately at each of the 12 stimulus orientations tested (see methods). The similarity between two decoders was quantified as the cosine of the angle between the decoder axes. For the contour stimuli, orthogonality was observed between the feature and object decoders across stimulus orientations, as manifested by a cosine similarity around zero (Fig. 5B, left, two off-diagonal quadrants; Fig. 5C, middle data point, not significantly different from zero, *P* = 0.12). In contrast, the object decoders at different stimulus orientations shared a similar axis (Fig. 5B, upper right quadrant; Fig. 5C, right data point, significantly different from zero, *P* < 10^−26^). Qualitatively similar results were observed for the other visual stimuli (singleton and surface in Fig. 5, B and C; streaked surface in fig. S8, B and C).

**Fig. 5.**
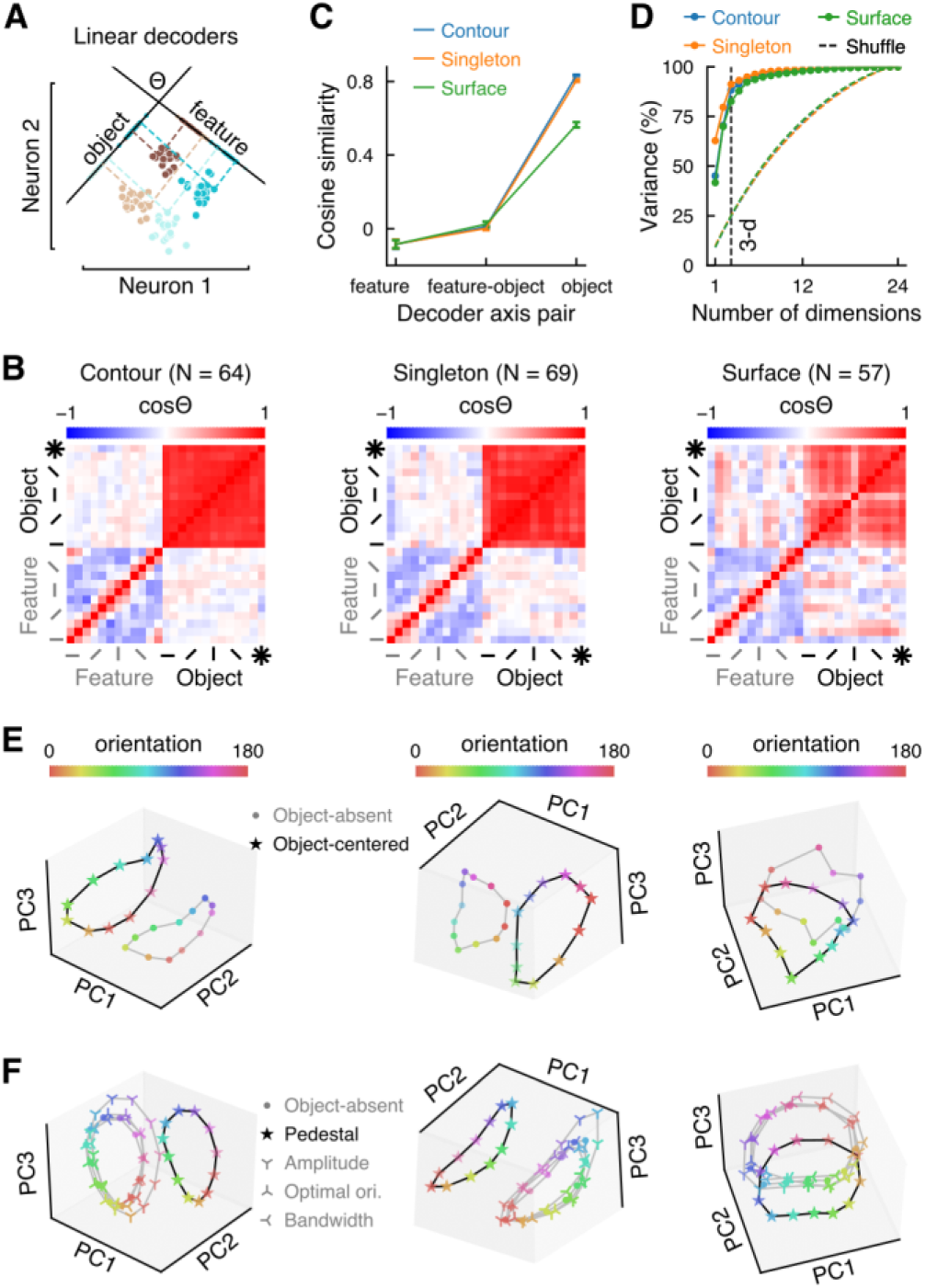
A convergent geometry of the segmentation manifold. The same dataset in Fig. 2 was used, but only orientation-selective V1 sites were included here. **(A)** Schematic feature and object decoders (see methods). Based on the trial-by-trial spike counts of individual neurons in response to a given stimulus orientation (brown) and its adjacent orientation (cyan) in the object-absent (light color) and object-centered (dark color) conditions, we trained separate FLD classifiers to discriminate the two stimulus orientations (feature decoder) and to differentiate the object-centered and object-absent conditions (object decoder). **(B)** Cosine similarity matrix of feature and object decoders for the contour (left), singleton (middle) and surface (right) stimuli, respectively. A cosine value of 0 (white color) indicates two orthogonal axes of the decoders to be compared. Combinations of all feature and object decoders at 12 stimulus orientations were compared (see methods). The object decoder marked with an asterisk was trained after pooling data from all stimulus orientations. **(C)** Mean cosine similarities computed from the matrices shown in B by pooling different stimulus orientations for the three types of stimuli, respectively. Shown are similarities between pairs of feature decoders (left), pairs of object decoders (right), and pairs of feature and object decoders (middle). **(D)** Fraction of explained variance as a function of number of dimensions obtained from PCA (see methods). **(E)** Orthogonal displacement of the ring-shaped orientation manifold (color coded) before (round dots) and after (stars) introducing an object (contour, singleton, or surface, respectively) into the background. **(F)** Similar to E, but showing changes in the neural manifold caused by respective changes in individual parameters (pedestal, amplitude, optimal orientation, and bandwidth) of the Gaussian fitted orientation tuning function in the object-centered relative to the object-absent condition. An increase in the pedestal (i.e., an additive elevation of the tuning curve) was the sole factor leading to an orthogonal displacement of the neural manifold (see methods for details). Note that similar results were obtained after pooling both orientation-selective and nonselective V1 sites (see fig. S7), and that the results from the streaked surface dataset were also consistent (see fig. S8).

Next, we sought to visualize the orthogonal geometry of the neural manifolds encoding bar orientation and the foreground object. Principal component analyses (PCA) revealed that the datasets were mostly three-dimensional (Fig. 5D and fig. S8D; 85-91% of variance explained in the raw datasets vs. ∼25% after trial shuffling). Geometrically, the bar orientation formed a 2-D ring-shaped manifold in the object-absent condition (Fig. 5E, fig. S8E). The presence of an object (contour, singleton, or surface) displaced the ring-shaped manifold along a single dimension orthogonal to the ring. Such an orthogonal geometry was well explained by the increased pedestal of the orientation-tuning curves rather than changes in other tuning parameters (Fig. 5F, fig. S8F), confirming the additive code in the population space.

## Computational logic of the segmentation code

How does the additive code in an early cortical area (e.g.,V1) affect downstream representations? We addressed this question by leveraging the convolutional neural network (CNN; Fig. 6A), which has been used as an *in-silico* model of the ventral visual pathway (*32, 33*). We simulated the impact of the additive code by adding activity to the layer of the CNN that best matched the cortical area V1 (the fourth layer L4; see fig. S9 and methods for layer selection). The additive code improved model performance when natural scene images (*34*) with different degradation levels (Fig. 6B) were fed into the CNN. The maximal improvement was seen at a degradation level of 30%, boosting the performance by 35% relative to the vanilla CNN. These results established an *in-silico* model for further simulating and dissecting the segmentation codes.

**Fig. 6.**
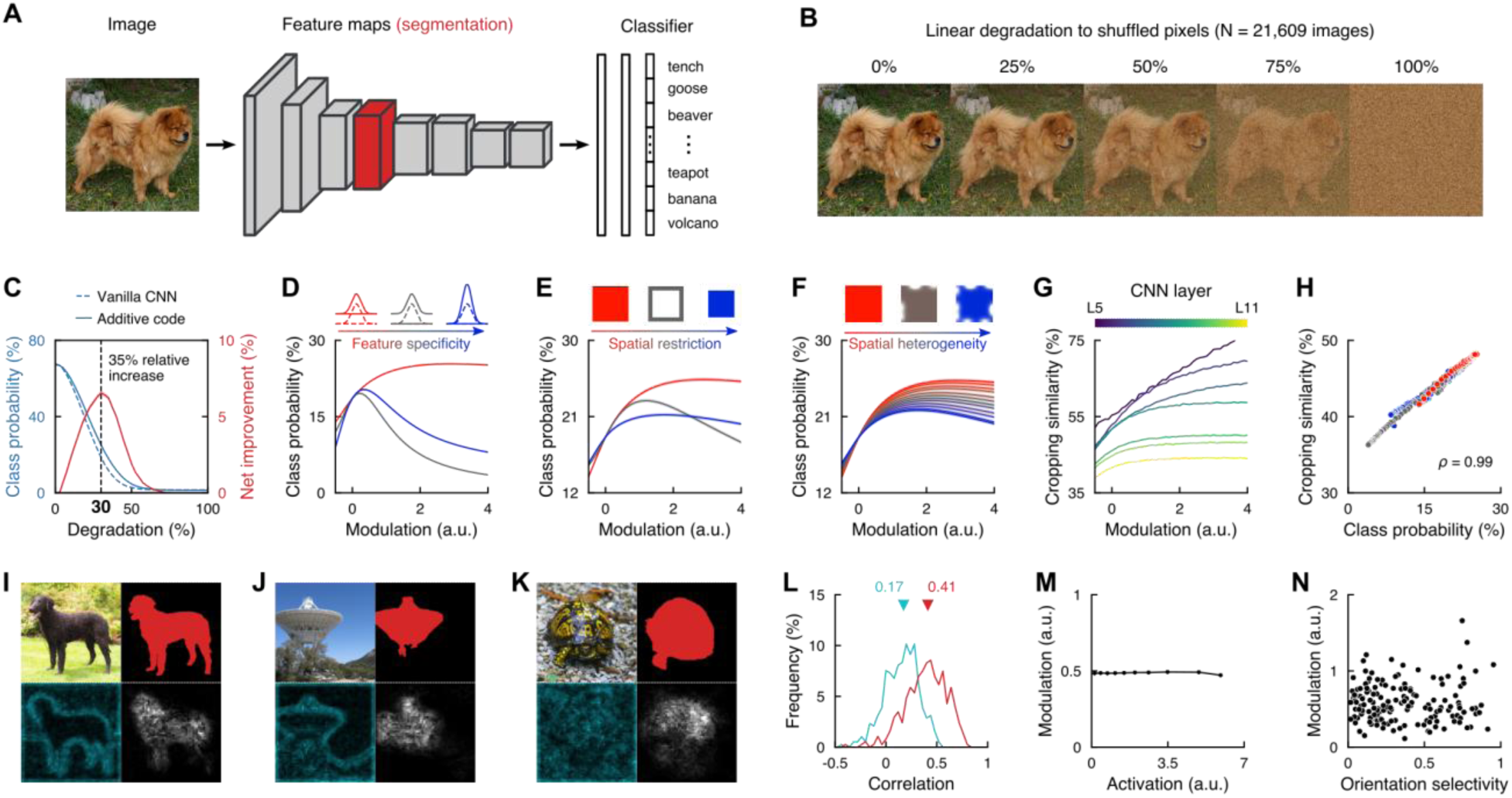
Computational logic of the segmentation code. **(A)** Architecture of the CNN used to model the impacts of different forms of segmentation codes on image classification. The fourth layer (L4, red) indicates the chosen V1 counterpart for implementing the segmentation code (see fig. S9 and methods). **(B)** A sample image and the linearly degraded variants (see methods). **(C)** Model performance on object classification, before (dashed blue) and after (solid blue) the inclusion of the additive code in L4, as a function of the image degradation level. The difference between the two curves indicates the net improvement caused by the additive code (red, right y-axis). Color shades indicate SEMs (barely visible). **(D)** Effects of different hypothetical segmentation codes added to L4 on the model performance examined at a range of modulation levels (see methods). Compared here are the three forms of codes with increasing specificity to oriented cues (see Fig. 1E). **(E)** Effects of introducing spatial restrictions to the additive code in L4 on the model performance (see methods). The biologically observed object-level code (red; as in D) is compared with the boundary-specific code (gray) and interior-specific code (blue). **(F)** Effects of locally attenuated additive code in L4 on the model performance. The additive modulations at a number of randomly-chosen boundary locations (0 to 11, color coded) were attenuated (see top insets) by a 2D Gaussian weight matrix (see methods). **(G)** Simulation test for the cropping hypothesis. For each layer downstream L4 (L5-11, color coded), the representational similarity was computed between two conditions: the intact images with the inclusion of the additive code in L4, and the background-cropped images (i.e., object region only) without introducing the code. Other hypotheses were also tested and excluded (see fig. S10, A and B). **(H)** Strong linear prediction of the model classification performance by the representational similarity to the cropped images. Here the representational similarity in the last fully connected layer (computed as in G) was compared with the model performance across all of the candidate codes and modulation levels (marked with different colors as in D to F). **(I** to **K)** Three examples of evolved modulations (see methods; see also fig. S10C). Top left quadrant: original image; top right: ground-truth object mask; bottom: activation (left) and modulation (right) maps of L4 units. **(L)** Distribution (across all the images tested) of the Pearson correlations between the evolved modulation map and the activation map in L4 (cyan), and between the modulation map and the object mask (red). **(M)** Independence between the evolved modulation and activation of L4 units. **(N)** Independence between the evolved modulation and the orientation selectivity of individual L4 units. The analyses in G to N were based on 1,838 images assembled from a subset of the ImageNet-S919 used for the simulations shown in B to F (see methods).

We first revealed the importance of the additivity and uniformity of the biologically observed segmentation code. By additively and uniformly modulating the baseline activations of L4 units in response to the object region, the model performance monotonically improved with increasing the modulation strength (Fig. 6D, red). However, once the segmentation modulation deviated from the additivity and uniformity, the model performance was generally decreased (Fig. 6D, gray and blue, introducing different degrees of feature specificity into the segmentation code; Fig. 6, E and F, introducing different forms of spatial restriction or heterogeneity).

Next, we examined how the additive code improves the model performance via influencing downstream representations. Among different hypotheses (see Fig. 6G and fig. S10, A and B), our simulations supported a cropping hypothesis: all downstream representations became more similar to the representations of isolated, background-cropped objects with increasing the additive modulation in L4 (Fig. 6G). Such a representational similarity seen in the last fully connected layer not only linearly predicted the performance improvement but also explicitly illustrated the advantage of the additive code over other candidate codes (Fig. 6H; *ρ* = 0.99, *P* < 10^−70^). Therefore, image segmentation in a V1-like layer improved the model performance by computationally extracting objects from the cluttered background, allowing downstream representations to focus on the segmented object.

Is the additive code a unique and optimal solution for image segmentation? The answer to this question is important considering the huge parameter space of the CNN. We trained the weight-fixed CNN to approximate downstream representations of isolated objects by optimizing a modulation matrix attached to L4 units. With training, the modulation matrix transformed gradually from a random state to a structured one that approximated the segmentation mask (Fig. 6, I to K, bottom right vs. top right; see also fig. S10C). This similarity was larger than the similarity between the modulation matrix and the baseline activations of L4 units (Fig. 6, I to K, bottom right vs. bottom left; Fig. 6 L, red vs. cyan, *P* < 10^−42^). Moreover, the evolved modulations depended neither on the baseline activations (Fig. 6M, *P* = 0.54) nor on the orientation selectivities of L4 units (Fig. 6N, *P* = 0.67), indicating an additive code directly evolved from mimicking downstream representations of isolated objects. These results argue that the biological additive segmentation code is indeed an optimal solution and that it is computationally equivalent to building a background-free object representation.

## Discussion

We found that distinct segmentation computations actually converge onto the same coding scheme: an object-level additive code in the earliest visual cortical area V1. This code generates a spatial segmentation mask for the entire object by uniformly enhancing V1 responses to the segregated object. It also translates the neural manifold for image features along an orthogonal dimension, efficiently multiplexing information about the local features and global saliency of the object in the same population of neurons. This coding scheme allows downstream analyses to focus on the segmented region and in turn unclutters object representations in cluttered environments.

### A spatial map of segmentation in V1

As a key intermediate step towards structured understanding of natural scenes, visual segmentation has been extensively studied from the behavioral perspective for over a century (*5, 6, 8*). After decades of physiological studies (*35–38*), we still lack characterization of several core issues.

First, a unified understanding of different segmentation computations is missing. Previous studies approach image parsing from different perspectives using assorted stimuli (*14, 16, 23, 31*). The current study, however, took a holistic approach and revealed a convergent cortical scheme across distinct segmentation computations.

Second, a systematic search for the determinants of the neural segmentation code is unavailable. By doing so, we found that the strength of the additive code reflects the saliency of the entire object (Figs. 2 to 4). The outcome is a uniform increase in V1 responses to the object, regardless of neuronal feature selectivities, RF locations on the object, and the local saliency around the RF. These characteristics are inconsistent with feature-based accounts of object segmentation and argue for a convergent spatial segmentation scheme, which likely is implemented in the late phase of image parsing (*14, 39*). This convergent scheme creates a spatial mask in V1 for object segmentation by overlaying the object-level saliency on top of local image features (*11*). Such a segmentation mask represents the ultimate product of diverse segmentation computations (*34*). In fact, segmentation masks have proven crucial in the field of computer vision and have been explicitly computed for a broad range of applications including object cropping (*40*).

Lastly, our results imply that the neural segmentation mask seen in V1 provides a critical link between low- and high-level visual representations (*2, 3, 7, 12*). As a prerequisite for such a linking role, the V1 codes for local features and for object segmentation are independent of each other, allowing for a faithful access to low-level features while building up object representations in downstream areas. The object mask implemented in V1 enables transformation of downstream object representations into uncluttered states. The linkage between low- and high-level vision provided by V1 represents a unified framework of the computational logic for image segmentation.

### An object-level code in V1

Our results revealed the crucial role of the object-level segmentation in V1, as manifested by the uniform highlighting of the object and its transformative effect on downstream object representations. The discovery of the object-level code in V1 argues for a revision of the current understanding of the visual cortical hierarchy, which posits a bottom-up, progressive transition from local to object-level representations (*41, 42*). Although an ascending gradient of feature complexities still holds in terms of neurons’ tuning properties, the earliest cortical stage is capable of interpreting local features in the context of the entire object. These results suggest that all of the hierarchically organized cortices could operate at the object level, with different areas specialized in different computational objectives. In this framework, V1 and inferotemporal cortex specialize, respectively, in segmentation and feature analysis of visual objects, collectively building an uncluttered object representation in cluttered environments.

The object-level segmentation code in V1 could also serve as a prop for other perceptual and cognitive processes. In line with the conjecture, selective attention can enhance V1 activities in an object-based manner by spreading additive modulation conforming to the Gestalt criteria (*16, 43, 44*). In addition, the segmentation code in V1 could also aid in binding of visual features belonging together, another process concerning visual representations in object-based manner (*45*). In this respect, the segmentation code could serve as an object pointer whereby selective attention can further highlight all features belonging to the same object.

## Acknowledgments

We thank Yin Yan and Xibin Xu for technical assistance and also thank Zhaoping Li and Tai Sing Lee for valuable comments.

## Author contributions

Conceptualization: MC

Methodology: MC

Investigation: MC, HC, WL

Visualization: MC

Funding acquisition: WL

Project administration: MC

Supervision: MC, WL

Writing – original draft: MC

Writing – review & editing: MC, WL, PRR

