## Supplementary material for "A convergent cortical scheme for visual scene segmentation": methods

### Materials and Methods

Ethical approval was granted by the Institutional Animal Care and Use Committee of Beijing Normal University, with all procedures in compliance with the National Institutes of Health Guide for the Care and Use of Laboratory Animals.

#### *Experimental model preparation*

**Subjects**

Two adult male monkeys (*Macaca Mulatta*, 6.5-7.3 kg) participated in this study. They were first trained in a simple fixation task by responding to a luminance change of a small point in the CRT center. After implantation of a head restraint (see below), they were further trained to maintain the gaze for a couple of seconds within an invisible window of 0.5° in radius around the fixation point. Eye positions were monitored by an infrared tracking system at 30 Hz (*46*).

**Surgeries**

All surgeries were conducted under general anesthesia induced with ketamine (10 mg/kg, intramuscular) and maintained, after intubation, by ventilation with $O_2$ (100%) mixed with isoflurane (1-2.5%). Vital signs such as $SpO_2$, $CO_2$, ECG and heart rates were monitored by a patient monitor (PM-9000 Express, Mindray) during the surgery. Antibiotics and analgesics were administered during recovery.

A biocompatible titanium head restraint was attached to the animal's skull with titanium bone screws under aseptic conditions. Microelectrode arrays (Utah Array, Blackrock Microsystems) were implanted using a pneumatic inserter (Blackrock Microsystems) into the dorsolateral portion of V1.

**Electrophysiological recordings**

Neuronal spikes were recorded using the implanted microelectrode arrays and a 128-channel data acquisition system (Blackrock Microsystems). Each array contained 6 × 8 electrodes (~0.5 mm in length spaced 0.4 mm apart). We implanted one array in one of the two animals and two arrays in the other. Spikes were detected by applying a voltage threshold with a signal-to-noise ratio of 3.5, and spike waveforms were sampled at 30 kHz.

The recorded spikes from an electrode (referred to as a V1 recording site) were sorted into clusters using customized software, whereby the informative multi-wavelet coefficients were extracted from the spike waveforms (*47*) and the spikes were then clustered by an Expectation-Maximization algorithm based on mixtures of multivariate *t*-distributions (*48*). This sorting procedure largely removed the noise due to random fluctuations of the local field potentials and separated the spikes from the same V1 site into different clusters. The cluster of spikes having the largest signal-to-noise ratio, the ratio between the mean peak-to-valley amplitude and the standard deviation of the waveform residuals, were taken as the spiking activity of the recorded site. Recording sites without a significant response to visual stimuli were excluded (see ***Data analyses***).

**Receptive field mapping**

While the animal was fixating, we measured the basic response properties of all the recording sites using square-wave gratings (1.5 cycles/deg, drifting at 2 cycles/s, 99% Michelson contrast with mean luminance of 21.2 cd/m$^2$). To map the receptive field (RF) positions along the horizontal axis, we used a vertical band of gratings (0.3° × 7°). Within a trial the location of the grating stimulus was fixed; only the gratings inside the band were drifting at 45° for 250 ms and then at 135° for another 250 ms. In different trials, the horizontal location of the grating stimulus was changed randomly in multiples of 0.3°, covering the RFs of all recorded sites. The RF positions along the vertical axis were mapped similarly by positioning a horizontal band of gratings (7° ×

0.3°) at different vertical locations. We also mapped the orientation selectivity of each V1 site using a large, circular grating patch (7° in diameter, 500 ms), with the grating orientation varied randomly in different trials. Within a block of trials each stimulus condition was repeated 5-10 times.

For each V1 site, its mean firing rates during stimulus presentation (0-500 ms) were plotted as a function of the grating positions or orientations. The resulting tuning curve was fitted with a Gaussian function (see ***Data analyses***). The horizontal and vertical positions of the RF, and the optimal orientation of an orientation-selective site, were defined as the corresponding Gaussian means. Based on the two Gaussians fitted respectively to the horizontal and vertical position tuning curves of a V1 site, the RF width or length was defined as 2 × 1.96 standard deviation (SD) of the corresponding Gaussian, subtracting the width of the grating stimulus (0.3°, see above) used for RF mapping. The recorded RFs from the two animals had mean eccentricities of 5.40° ± 0.34° (mean ± SD) and 4.67° ± 0.72°, respectively; mean widths of 0.83° ± 0.18° and 0.67° ± 0.23°; and mean lengths of 0.70° ± 0.19° and 0.65° ± 0.19°.

***Experiments***

**Visual stimuli and behavioral paradigms**

Visual stimuli, anti-aliased and viewed at a distance of 100 cm, were generated by a visual stimulator (ViSaGe MKII, Cambridge Research Systems) on a gamma-corrected CRT monitor (Vision Master Pro-514, Iiyama; 40.4 × 28.8 cm with a resolution of 1,200 × 900 pixels at 100 Hz).

The stimuli used for 3 different segmentation scenarios (Fig. 1F) were all composed of short bars (0.25° × 0.05°) distributed in a hidden grid of 0.5° squares, each containing a bar with a random position jitter of up to 0.075° from the square center. The constituent bars were white (12.4 cd/m$^2$) on a uniform gray (4.1 cd/m$^2$) background. For the contour stimulus (Fig. 1F, left), a number of adjacent bars in a row were collinearly aligned and evenly spaced, leaving the remaining bars at random orientations and jittered locations. The saliency of the object (i.e., the contour) could be controlled by varying the number (1 to 9) of collinear bars in the display (fig. S1A). For the singleton (Fig. 1F, middle) and surface (Fig. 1F, right) stimuli, the bars belonging to the object (1 bar for the singleton; 9 × 9 iso-oriented bars for the upright square surface) were arranged at an orientation different from the remaining iso-oriented bars belonging to the background. The saliency of the object could be controlled by adjusting the angular difference (0-90°, termed the orientation contrast; fig. S1, B and C) between the object and background bars.

In a two-alternative forced-choice task (fig. S2), the animal was required to indicate where the simple object (contour, singleton, or surface) was located in the display. For these three segmentation scenarios, there was a difference in the arrangement of stimulus display. For the singleton and surface stimuli (fig. S2, B and C), the entire display was occupied by the background bars except that the object (singleton or surface) was randomly embedded at one of two predefined locations that were symmetrical about the fixation point: One location coincided with the neuronal RF to be examined in the lower visual field; the other was in the opposite, upper visual field. For the contour stimulus (fig. S2A), the stimulus display in a trial was composed of two circular stimulus patterns (9.5° in diameter) centered respectively at the two predefined locations; and the contour could appear at either location with equal probability. A trial began when the monkey fixated within an invisible circular window (0.5° in radius) around the fixation point. After 300 ms of fixation, the stimulus appeared for 500 ms, in which an object (contour, singleton, or surface) was embedded in the background randomly at one of the two predefined locations (see above).

Subsequently the stimulus disappeared, and two dots were presented at the two locations. The monkey was rewarded for making a saccade within 800 ms to the dot corresponding to the location where the object had been shown. Trials were aborted if the animal's gaze shifted out of the fixation window during stimulus presentation. Within a block of trials, only one type of stimulus (contour, singleton, or surface) was used.

Another popular and more complex design of surface segregation (*23*) was also examined for comparison of neuronal responses (fig. S4A, referred to as the streaked surface). In this particular case, two 4° upright texture squares were centered at the two predefined locations (see above) and surrounded by a full-screen texture background. The streaked textures within each of the three designated areas (the two squares and the entire background) were generated independently. On top of each area, an invisible grid with 0.15° spacing was randomly positioned. Each node of the grid was assigned a 0.48° × 0.03° line segment with a position jitter up to 0.3°. The part of the line segment falling outside the designated area was cropped. All line segments were dark (4.1 cd/m$^2$) on a brighter (12.4 cd/m$^2$) uniform background. The orientation of line segments forming one texture square (the object) was always set orthogonal to the line segments forming the full-screen texture background and the other texture square (the sham object).

**Behavioral experiments**

We measured the animal's behavioral performance on segmenting the object from the background at different levels of object saliency. The contour saliency (fig. S1A) was controlled by aligning 1, 3, 5, 7 or 9 collinear bars. Note that in the 1-bar contour condition, the two circular stimulus patterns on the screen (see fig. S2A) were identical and contained no object (*49*). The saliency of singleton (fig. S1B) and surface (fig. S1C) were controlled by adjusting the orientation contrast between the object and background bars, from 0° (i.e., no object) to 90° in 15° steps. Stimulus conditions with the same type of stimuli (contours, singletons, or surfaces) but different object saliencies and two possible object locations were randomly interleaved in a block of trials; and each condition was repeated in at least 300 trials for a reliable estimation of the behavioral accuracy. In each trial, the component bars of the object were set at one of 12 orientations (0° to 180° evenly spaced) in a random manner (see **Orientation independency experiments** below for details on orientation manipulation). For a direct comparison across different segmentation scenarios, the saliency levels of different objects were rescaled within the range of 0 to 1 (Fig. 1G; behavioral results at the original non-scaled saliency levels were presented in the captions of fig. S1).

**Orientation independency experiments**

To examine whether the neural code for image segmentation was independent of neurons' orientation selectivity, the orientation tuning curve of a V1 site was sampled from 0° to 180° in 15° steps when the object was placed respectively at the two predefined locations (see **Visual stimuli and behavioral paradigms**). Each orientation and object location were repeated in at least 30 trials. When the object was centered at the RF location, we termed it the object-centered condition (fig. S3, A, F, K, and fig. S4A, left column); when the object was located in the opposite visual field and the RF was merely covered by the background stimuli, we termed it the object-absent condition (fig. S3, A, F, K, and fig. S4A, right column). Specifically, for the contour stimuli containing a 9-bar contour (fig. S3A), when the recorded RF was centered on the middle component bar of the contour in the object-centered condition, or on a background bar in the object-absent condition, we measured neuronal orientation tuning by rotating the two circular stimulus patterns in a trial (see fig. S2A) around their respective centers. For the singleton and surface stimuli (fig. S3, F and K) and the streaked surface (fig. S4A) having a 90° orientation contrast, we varied the orientations of individual bars forming the object and background by the same amount in both the object-centered and object-absent conditions, keeping the object outline and saliency unchanged.

**Location independency and object-level saliency experiments**

In the above object-centered conditions, the RF of a V1 site was always centered on the middle component bar of the object. To test the conjecture of object-level saliency that the neural code for image segmentation could be independent of the RF location on the object, the component bars at different distances from the object center were respectively placed in the RF, varying the relative location of the RF on the object (valid for the contour and surface but not the singleton; Fig. 3, A and D). Only orientation-selective V1 sites were included; and for each site, the bars forming the object (contour or surface) were optimally oriented. Each RF location was tested in at least 30 trials.

To further test the conjecture of object-level saliency, we introduced heterogeneity to the component bars forming the global contour and surface, as described below.

To construct a contour with heterogeneous elements (Fig. 3G, top), we started with two short contour fragments having different local saliencies: a discontinuous fragment consisting of two collinear bars interrupted in between by an orthogonal bar (Fig. 3G, middle; termed the triplet here); and a continuous fragment of two adjacent collinear bars (Fig. 3G, bottom). These two contour fragments were merged along a linear path into a longer, central segment, which was further elongated by appending to each side an additional contour segment with 4 interdigitated bars: 2 randomly oriented and 2 collinear with the contour path (Fig. 3G, top). This manipulation created the percept of a global object, which we termed the chimera contour. We measured neuronal responses to the chimera contour by centering the RF respectively on each component bar in the triplet (see above). We balanced and pooled two complementary sets of stimuli (fig. S5A, top row): The contour path was either aligned with, or orthogonal to, the optimal orientation of the recorded V1 site; so was the component bar in the RF. As a control, we also tested the condition that all component bars of the chimera contour were collinearly aligned (fig. S5A, bottom row).

Similarly, based on a square surface formed by 9 × 9 bars with 90° orientation contrast against the background bars, a chimera surface (Fig. 3K, top) was created by gradually morphing the orientation contrast of a subset of the surface bars from 90° (Fig. 3K, bottom) to 20° (Fig. 3K, middle). The morphing was restricted to a semicircular surface region whose diameter coincided with one boundary of the square surface (Fig. 3K, top). For a bar within this region, its orientation was altered clockwise by a specific amount (between 0 and 70°) proportional to its distance from the invisible semi-circumference. As a result, the surface bars along the semi-circumference remained unaltered, keeping their original orientation contrast of 90°; but the surface bar located at the diameter center was rotated by the maximum amount of 70°, thus having the least orientation contrast of 20° relative to the background bars. The RF of a V1 site was centered either on this particular bar (Fig. 3K, top, green circle), or on the corresponding bar at the opposite boundary of the chimera surface where the orientation contrast was 90° (Fig. 3K, top, purple circle). For each of the two cases, two complementary stimulus sets were balanced and pooled: The surface component bar in the RF was optimally or orthogonally oriented, keeping its orientation contrast unchanged relative to the background (fig. S5B, top and middle rows). As a control, the singleton stimulus was also tested similarly (fig. S5B, bottom row).

**Dissociation of the effects of local and global orientations**

This experiment was designed to dissociate the effects of local orientation (i.e., a component bar) and global orientation (i.e., the object as a whole) by focusing on orientation-selective V1 sites.

To examine the object-level orientation selectivity of a V1 site for the global contour, we set the contour path either at the optimal orientation of the V1 site or at the orthogonal orientation (Fig. 4A). The contour consisted of collinear bars except for the middle component bar, which was centered in the RF and kept at 45° relative to the optimal or orthogonal orientation regardless of the global contour orientations. Similarly, we measured neuronal selectivity for the local orientation of the middle component bar by setting it at either the optimal or orthogonal orientation, keeping the contour orientation 45° in between (fig. S6A). Note that the saliency of the contour was exactly identical when the global or local orientation was varied.

For the surface stimulus, a slim version (a single row of iso-oriented bars among a background of orthogonal bars; Fig. 4D) was used to probe the orientation selectivity of a V1 site for the elongated surface (global orientation) and for its middle component bar (local orientation), respectively. The global orientation was varied simply by rearranging the orientation of the hidden grid used for placing the bars in the stimulus (see **Visual stimuli and behavioral paradigms**). Specifically, in a block of trials, we randomly set the grid rows at one of 12 orientations (0° to 180°, 15° apart). The iso-oriented bars forming the slim surface were arranged along a grid row to define the global orientation; and the local orientation of surface bars was also randomly set at one of the 12 orientations. The background bars were always kept orthogonal to the surface bars to maintain a constant orientation contrast. By pooling the 12 local bar orientations for each surface orientation, or by pooling the 12 surface orientations for each local bar orientation, the global (Fig. 4E) and local (fig. S6E) orientation tuning curves were dissociated. The resulting global or local tuning curve in the object-centered condition was compared with the corresponding curve constructed similarly in the object-absent condition.

In the above experiments, each of the global and local orientations was tested in at least 30 trials.

**Multiple objects experiments**

To examine whether multiple objects were segmented independently by V1 neurons in a simple additive manner, we compared neuronal responses in 3 stimulus conditions, in which 0, 1, and 2 objects were embedded in the same background (Fig. 4, G and J for the contour and slim surface, respectively). The stimulus display in the absence of any embedded object was used as a baseline condition (Fig. 4, G and J, top row; object-absent). In the presence of one object, we measured neuronal responses in two conditions: The object was either centered on the RF (Fig. 4, G and J, 2nd row; object-centered), or placed 1° away from the RF center where a background bar was located (Fig. 4, G and J, 3rd row; object-flanked). By introducing another parallel object to the RF location in the object-flanked condition, neuronal responses were further examined in the presence of two objects (Fig. 4, G and J, bottom row; doublet object).

In addition to the firing rates, we also investigated the spike-spike coherence between a pair of V1 sites when their RFs were located on the same object or different ones (Fig. 4, M and P). We focused on pairs of sites with RFs separated by 0.6° to 1.5° (center to center) and with optimal orientations differed less than 30°. With these settings, the synchronization effect, if any, would be maximized (*50, 51*). The size of the invisible grid used for stimulus generation (see **Visual stimuli and behavioral paradigms**) was adjusted so that a pair of recorded RFs could be centered on the designated component bars of the objects. All stimulus conditions were repeated in at least 40 trials for a reliable estimation of the coherence.

***Data analyses***

**General analyses**

Spearman's rank correlation was used to quantify the relationship between object saliency and behavioral accuracy. Statistical significance of the correlation coefficient was determined based on its Student's $t$ approximation (*52*).

Only recording sites showing significant responses to any of the stimulus conditions were included in the analysis; statistical significance was determined at the level of 5% using the Wilcoxon signed-rank test by comparing the mean stimulus-evoked activity (0-500 ms relative to stimulus onset) with the mean spontaneous activity (−300 to 0 ms). Each site was included only once in each analysis; and each pair of sites was also included only once in the inter-neuronal correlation analysis. The V1 sites from the two animals in the same experiment were pooled as the results were qualitatively similar (see the scatter plots in Figs. 2 to 4, figs. S4 and S6, where light and dark data points represent individual V1 sites from the two animals, respectively).

The peri-stimulus time histogram (PSTH; e.g., Fig. 2, A, G and M) for each recording site under a stimulus condition was created by binning the spikes in 1 ms intervals and averaging the binned spike counts across trials. The raw PSTH was smoothed by convolving it with a Gaussian window (SD = 5 ms). The PSTHs of each V1 site in all the stimulus conditions to be compared were normalized to the largest peak of these PSTHs, with the mean spontaneous activity subtracted. The normalized PSTHs for a given condition were averaged across V1 sites to derive the population PSTH. The above normalization procedure scaled neural responses so that the maximum was unity (indicated as a.u. in all related figures).

Except for the RF mapping, analyses of mean firing rates were based on the late components of V1 responses (100-550 ms) containing marked segmentation signals (*14, 16, 24*). We defined the segmentation signal as the difference of normalized mean V1 responses between the paired object-centered and object-absent conditions.

**Quantification of neuronal orientation selectivity**

Based on neuronal responses to the gratings of different orientations (see **Receptive field mapping**), V1 sites were classified into the orientation selective or nonselective groups by two statistical tests. For each site, we first used the Kruskal-Wallis test to examine whether the grating orientation was a significant influencing factor. We then performed a permutation test to determine the significance of a Gaussian fit to the tuning curve r(θ):

$$r(\theta) = p + ae^{\frac{-(\theta-\mu)^2}{2\sigma^2}} \quad \text{(Equation 1)}$$

where p, a, μ, and σ stands for the pedestal, amplitude, optimal orientation, and bandwidth of the tuning function. A cross-validation procedure was employed to minimize overfitting due to limited sampling of stimulus orientations. Specifically, the trials for each stimulus orientation were randomly split into two halves. One half was used for estimation of the Gaussian parameters, based on which the remaining half was used for computing the goodness of fit ($R^2$). To test for statistical significance of the $R^2$ we used a permutation approach, whereby individual trials were randomly assigned to one of the stimulus orientations, and the above cross-validation procedure was performed on the permuted dataset. After 1,000 repetitions of this process, the *P* value was computed by comparing the $R^2$ derived from the raw dataset against the distribution of $R^2$ values from the permuted ones. A V1 site was classified as orientation selective only if both of the following criteria were met: $P < 0.05$ in the aforementioned Kruskal-Wallis test for orientation dependence of neural responses; and $P < 0.05$ in the permutation test for significance of the $R^2$ of the Gaussian fit to the original tuning curve. Otherwise, the V1 site was regarded as orientation nonselective.

For each recording site, based on its mean responses to the gratings placed at the orthogonal and optimal orientations, we defined its orientation selectivity as the ratio of the two mean responses (orthogonal/optimal after subtracting the mean spontaneous activity; commonly referred to as the OP ratio) (*53*).

Orientation tuning curves were also measured in the segmentation experiments (see **Orientation independency experiments**; **Dissociation of the effects of local and global orientations**). To construct the population tuning curve averaged across V1 sites, individual curves from the orientation-selective sites were aligned to their respective optimal orientations (denoted as the relative orientation 0 in Fig. 2, C, I, O, and fig. S4H; Fig. 4E and fig. S6E). For the nonselective sites, a random orientation was assigned to each site as the optimal before averaging over the population.

Based on the paired Gaussians fitted to the orientation tuning curves in the object-absent and object-centered conditions, we compared the changes in each of the 4 tuning parameters (Equation 1) for orientation selective sites (Fig. 2, D, J, P, E, K, and Q; fig. S4, I and J; Wilcoxon signed-rank test for statistical significance of the changes). The respective contributions of the 4 parameters to the overall changes in the tuning curve were dissociated by taking the following approach. To estimate the contribution of a parameter $c$ ($c$ denotes any of the 4 parameters in Equation 1), we first constructed a surrogate Gaussian tuning curve for a given V1 site by combining the $c$ in the object-centered condition with the other 3 parameters in the object-absent condition. This surrogate Gaussian tuning curve isolated the *specific* tuning changes caused by the $c$ in the object-centered relative to the object-absent condition. We next computed the *overall* changes between the original paired Gaussian tuning curves in the object-centered relative to the object-absent condition. After averaging over all stimulus orientations for the above *specific* and *overall* changes, respectively, we took the ratio of the two average values as the proportion of tuning changes explained by the parameter *c* for the given V1 site. We then quantified the population-level contribution of the parameter *c* by averaging all the ratios derived from individual orientation-selective sites (Fig. 2, F, L and R; fig. S4K).

Pearson correlation was used to quantify the relationship between the segmentation signal (defined in **General analyses**) and the orientation selectivity (the OP ratio defined above) across individual V1 sites (Fig. 2, B, H and N; fig. S4G). Statistical significance of the correlation was determined by the t-test (*54*).

**Linear prediction of neural responses to multiple objects**

We recorded the responses of a V1 site in the presence of 0, 1, and 2 objects (see **Multiple objects experiments**; Fig. 4, G and J). Three types of neural signals were computed and compared. Neuronal responses in the object-absent condition were used as the baseline signal. In the presence of one object, the object-enhancement signal was computed by subtracting the object-absent condition from the object-centered condition; and the background-suppression signal with a negative value (*14*) was isolated by subtracting the object-absent condition from the object-flanked condition in which the object was laterally displaced by 1° from the RF center. With the insertion of the second object to the RF location flanked by the first one, we tested the linear prediction that the neuronal responses in this doublet object condition were simply the sum of the above 3 types of signals: the object-enhancement signal (induced by the centered object) and the background-suppression signal (induced by the flanked object) superimposed on the baseline signal (in the object-absent condition). The Wilcoxon signed-rank test was used to assess the difference between this linear prediction and the observed neuronal responses.

**Inter-neuronal correlation**

Neuronal interactions were examined by computing spike-spike coherence, a measure of similarity in each frequency component of spike trains between the paired V1 sites (see **Multiple objects experiments**). The time series of firing rates in a trial was constructed by binning the spikes in 1 ms intervals. The power spectra of the spike trains were estimated using the multitaper technique (*55*). Coherence between two signals was defined as the modulus of their cross-spectrum normalized by the geometric mean of autospectra (Fig. 4, N and Q). The coherence values between a pair of V1 sites were then averaged over all frequencies for statistical tests (Fig. 4, O and R).

**Population decoding**

By pooling the data from individual V1 sites recorded separately in different sessions of the **Orientation independency experiments** (Fig. 2; fig. S4), we trained linear classifiers (Fisher linear discriminant, FLD) to discriminate two adjacent stimulus orientations (15° apart) or to differentiate the object-centered and object-absent conditions (see the illustration in Fig. 5A).

For each V1 site, we randomly resampled the trials with replacement 1,000 times for each stimulus orientation in the object-centered and object-absent conditions, respectively. Based on the trial-by-trial spike counts of all the V1 sites, a FLD (named the feature decoder) was trained to discriminate between a given stimulus orientation $O_i$ ($i$ = 1 to 12 covering 0-180° range) and the adjacent, counter-clockwise orientation $O_{i+1}$ after pooling the object-centered and object-absent trials; and a separate FLD (named the object decoder) was trained to discriminate between the object-centered and object-absent conditions for the given $O_i$. By repeating the above steps 1,000 times, an average decoder axis was computed respectively for the feature and object decoder at $O_i$. We repeated this decoding procedure to cover all the 12 stimulus orientations ($O_1$ - $O_{12}$).

The cosine similarity index was then used to quantify the relationships between the feature and object decoder axes (Fig. 5, A and B; fig. S7, A and B; fig. S8, A and B). A cosine value of 0 indicates two decoder axes orthogonal to each other, implying independence of the neural codes for the visual feature (local bar orientations) and for the segmented object (contour, singleton, or surface).

**Manifold visualization**

To visualize the geometric structures in the population responses of neurons to a range of stimuli (i.e., the neural manifold), we first evaluated the dimensionality of V1 population responses in the **Orientation independency experiments** (Fig. 2; fig. S4). For each V1 site, its mean firing rates across trials were calculated respectively for the 12 orientations in the object-centered and object-absent conditions. The resulting 12 × 2 matrix of firing rates was flattened as a 24-D vector. We stacked the vectors from different V1 sites and performed the principal component analysis (PCA) on the stacked matrix. Since most of the variance of the dataset could be explained by the first three principal components (PCs; Fig. 5D, fig. S7D, and fig. S8D, solid curves), we visualized the neural manifold in a 3-D scatter plot based on the embeddings of the 24 conditions in the three PCs (Fig. 5E, fig. S7E, and fig. S8E).

To further confirm the low-dimensional nature of the V1 dataset, we shuffled the 12 × 2 condition labels of all trials and repeated the above PCA procedure. By averaging over 1,000 repetitions of the trial-shuffled analysis, we plotted the fraction of variance explained by different number of PCs (Fig. 5D, fig. S7D, and fig. S8D, dashed curves).

Similar to dissociating the respective contributions of the 4 orientation-tuning parameters (Equation 1) to the overall changes in the tuning curve (see **Quantification of neuronal orientation selectivity**), we also dissociated the respective contributions of the 4 parameters to the changes in the neural manifolds between the object-absent and object-centered conditions (Fig. 5F, fig. S7F, and fig. S8F). We first synthesized a surrogate dataset for each Gaussian parameter $c$ ($c$ denotes any of the 4 parameters in Equation 1) so that its contribution to the neural manifold changes could be isolated. Specifically, for a given parameter $c$, by combining its value in the object-centered condition with the other 3 parameters in the object-absent condition, a surrogate Gaussian tuning curve was produced for each V1 site. By pooling all recorded sites, a dataset of neuronal responses (12 stimulus orientations × N sites) was synthesized from the surrogate tuning curves. This synthesized dataset isolated the changes caused by $c$ in the population space between the object-centered and object-absent conditions. Following the same procedure, a total of 4 datasets were generated for the 4 Gaussian parameters. These datasets were concatenated, together with another surrogate dataset derived directly from Gaussian fitted tuning curves in the object-absent condition (also 12 × N). The PCA was then performed on the concatenated matrix (60 × N), and the neural manifold corresponding to the change in an individual tuning parameter was visualized based on the embeddings in the first three PCs (see Fig. 5F, fig. S7F, and fig. S8F).

***Visualization of computational schemes supporting image segmentation***

**Visualization of various computations in segmenting natural scene images (Fig. 1B)**

Although multiple complementary computations have been proposed for image segmentation (*9, 10, 16, 49*), their respective effects remain unclear. We sought to visualize the outputs from different segmentation computations for segmenting natural scene images.

By implementing the so-called association field (*10*), we simulated the impact of contour continuity (collinearity or co-circularity) (*12, 56*) on the output of edge filters (Fig. 1B, left). After convolving an image (Fig. 1A) with a pair of 3 × 3 horizontal and vertical Sobel edge filters (*57*), orientation angle and energy information was assigned to each pixel in the image. Next, an association field (AF) was generated respectively for each pixel, which defined the AF center and its base orientation; the remaining pixels in the image constituted the contextual pixels of the AF. Each AF contained 3 component maps: First, a continuity map for indicating—at each contextual pixel location—the ideal orientation strictly collinear or co-circular to the AF center; second, a continuity weight map defined as the cosine of the orientation difference (i.e., cosine similarity) between the AF center and each of the pixels on the continuity map; and third, a spatial weight map defined as an isotropic 2D Gaussian (amplitude = 1, SD = 5 pixels). For any given contextual pixel, its influence on the AF center was the product of 4 factors: its orientation energy in the image; the cosine similarity between its de facto orientation in the image and its ideal orientation on the continuity map; and the other two factors combining the weight values of the contextual pixel on the continuity weight map and the spatial weight map described above. The influences of all the contextual pixels on the AF center were summed, which simulated the impact of edge continuity on the pixel in the AF center. To simulate the overall visual effect of edge continuity in the entire image, we applied the above procedures to all the pixels and performed 15 iterations of the entire process (Fig. 1B, left).

Unlike detection of edge continuity, visualization of the effects of feature similarity and dissimilarity requires a variety of filters. We adopted those filters in the first convolutional layer of VGG11, a popular deep neural network for image processing (*58*). The set of filters, a total of 64 and each 3 × 3 in size, are capable of detecting not only the orientation energy but also other features such as complex patterns and colors. The image was convolved with these filters to generate a full representation of visual features at each pixel. Next, we computed the Pearson correlation between the representation vectors of each pair of pixels. The feature similarity (Fig. 1B, right) was quantified as the product of the Pearson correlation coefficient and an isotropic 2D Gaussian weight map (amplitude = 1, SD = 5 pixels). Likewise, the feature dissimilarity (Fig. 1B, middle) was quantified as the product of the Pearson correlation coefficient and an isotropic 2D weight map of DoG (difference of Gaussians; amplitude = 2 and −1, SD = 1.25 and 5 pixels, respectively for the two Gaussians).

In addition to the three complementary segmentation computations explored here (contour continuity, feature similarity and feature dissimilarity), it is reasonable to assume that image segmentation based on other regularities or disruption of regularities could also be quantified by applying appropriate filters, and that better segmentation effects could be achieved with more elaborate filters.

**Visualization of hypothetical segmentation codes (Fig. 1D** and **E)**

To demonstrate the visual effects of hypothetical segmentation codes, we focused on image segmentation in the orientation domain. Two orthogonal orientation-selective filters (Fig. 1A, 2nd panel, left column) and two orientation-nonselective filters (Fig. 1A, 2nd panel, right column) were used for the visualization. These filters were selected among those in the first convolutional layer of AlexNet (*59*) for their resemblance to V1 RFs with and without orientation selectivities. We first computed the baseline activations simply by convolving the entire image with the four filters, respectively (Fig. 1D). Three types of hypothetical segmentation codes were then visualized by modulation of the baseline activations in different manners (Fig. 1E; see details below). The modulation was restricted to the object region delimited by the ground truth object mask (*34*).

The coding scheme for image segmentation simply by orientation cues could in effect come in three possible forms with different degrees of specificity to the oriented cues (Fig. 1E). For an orientation-selective neuron with RF covering a bar belonging to the object, its response could be modulated in a multiplicative manner whereby the modulatory strength depends on the orientation of the bar in the RF: A maximal modulation is seen if the bar is optimally oriented, but no modulation at all if the bar is orthogonally oriented. In order to segment an image, this coding scheme specifically and differentially modulates the responses of neurons with different orientation selectivities, entirely ignoring nonselective neurons. Therefore, the modulatory patterns of neural activities will be highly dependent on the oriented cues contained in the image, showing strong stimulus specificity (Fig. 1E, left, high specificity). The effects of this highly specific coding scheme were visualized by a multiplicative, fivefold increase in the outputs from the two orientation-selective filters, respectively, in response to the object region, leaving the two nonselective filters untouched. The second candidate coding scheme shows less specificity to oriented cues (Fig. 1E, middle, medium specificity): Instead of differentially modulating orientation-selective neurons in a multiplicative manner, this coding scheme uniformly enhances the responses of all orientation-selective neurons with RFs on the object, regardless of their optimal orientations; but it does not modulate orientation-nonselective neurons. In other words, this code is restricted within the orientation domain, but it is insensitive to specific oriented cues. We visualized this code by a constant increment of 5 times the mean response in the outputs from each orientation-selective filter in response to the object region, keeping the outputs from the two nonselective filters unchanged as in the above highly specific scheme. The third candidate coding scheme without any specificity was termed the general additive code (Fig. 1E, right, no specificity; simply referred to as the additive code throughout the current study). This general additive scheme uniformly enhances responses of all neurons irrespective of their orientation tuning. We visualized its effects by a constant increment of 5 times the mean response in the outputs from each of the 4 filters.

***Computational logic of the segmentation code (Fig. 6)***

To better understand the inner workings of the segmentation code, we sought to establish an in-silico model by leveraging the convolutional neural network (CNN), which has proven effective for modeling the ventral pathway of the visual system (*60, 61*). Specifically, we focused our analysis on VGG11 (*58*), a variant in a CNN class whose middle layers effectively model the macaque V1 (*62*). Unlike the biological system, this version of CNN only consists of feedforward connections without any build-in recurrence. Although recent developments have incorporated various forms of recurrence (*63*), we preferred the feedforward version for straightforward implementation of hypothetical codes in this architecture.

Three image datasets were used for different purposes. One dataset consisted of simple gratings for probing orientation representations of CNN units. Full field images (224 × 224 pixels) were generated by varying the orientation of sine-wave gratings from 0° to 180° in 15° steps. For each orientation, we also varied the grating spatial frequency (1/200, 1/100, 1/50, 1/25, and 1/12.5 cycles/pixel) and phase (0° to 360° in 15° steps). The second dataset, known as ImageNet-S919 (*64*), consisted of natural scene images for studying the effects of different hypothetical segmentation codes on CNN's classification performance. Although this dataset contained only a subset of images (21,609) from the original ImageNet, it provided the ground truth segmentation mask for the object to be classified (*34*). The third dataset, which was a further reduced subset (1,838 out of 21,609 in the second dataset), was used to study the nature of changes in downstream representations caused by introducing the segmentation codes into the CNN (see **Transformation of downstream representations** below for details).

All the readouts and manipulations of CNN activities were targeted to the ReLU units attached to the convolutional and fully connected layers, as ReLU outputs are analogous to neuronal spike rates. Max pooling and normalization layers were skipped in the analysis, as an intuitive mapping of these layers onto the biological network is absent. The model performance was quantified as the softmax class probability from the CNN output.

**Layer selection**

We determined the CNN layer used for incorporating the segmentation codes according to three prerequisites. First, it should be a convolutional layer since image segmentation relies on grouping of visual features in the spatial domain. Second, its activities should resemble orientation representations in V1 (as shown in the orientation manifold, Fig. 5E). Third, an inclusion of the biological segmentation code in the selected layer should improve the model performance on segmenting images. We conducted the following simulations to select the CNN layer conforming to these criteria.

We compared the representations of grating orientations across different convolutional layers by presenting the grating dataset to the CNN. The mean response to each orientation was computed for every unit in each convolutional layer by averaging over all grating spatial frequencies and phases. We then conducted PCA of the units' responses and plotted the orientation manifold for each layer using the first two PC embeddings (fig. S9, A to H).

To examine the impact of the inclusion of the additive code on CNN's classification performance, unit activations in response to the object region were uniformly increased by the addition of 4 times the average response of all units in the same layer. The inclusion of the additive code was tested respectively for different convolutional layers. The model performance was evaluated by averaging the softmax class probability of individual images (fig. S9I).

Based on the above simulations, the fourth convolutional layer (termed L4) was chosen as the counterpart of monkey V1.

**Image degradation level selection**

The images in the ImageNet-S919 database usually have high signal-to-noise ratios, imposing a limit on exploitation of the effect of the segmentation code. To overcome this limitation, we progressively degraded the images, with the degradation level determined by a weighted sum of the original image and its pixel-shuffled version (Fig. 6B). A total of 41 degradation levels were tested, from 0% to 100% in 2.5% steps. The degradation level of 30% (i.e., the linear sum of 30% of the shuffled image and 70% of the original image) was chosen for subsequent analyses, because at this degradation level the largest performance boost was observed after the inclusion of the additive code in L4 (Fig. 6C).

**Implementation of the hypothetical codes in the CNN**

By leveraging the CNN, the influences on image classification performance were compared among the three types of hypothetical codes with different degrees of specificity to orientation cues (see **Visualization of hypothetical segmentation codes**).

To this end, we first analyzed the orientation selectivities of CNN units. The statistical method used for V1 neuron classification (**Quantification of neuronal orientation selectivity**) could not apply to artificial CNN units for two reasons: The statistical approach relies on trial-by-trial variations of neuronal activations by the same image, distinct from the deterministic nature of the CNN architectures and unit responses; the OP ratio quantifying neuronal orientation selectivity is not suited for the non-Gaussian orientation tuning shape and for weak grating activations of some

CNN units. Therefore, we used another popular measure of orientation selectivity, the circular variance (CV):

$$CV = 1 - \left|\frac{\sum_{k=1}^{12} r_k e^{i2\theta_k}}{\sum_{k=1}^{12} r_k}\right| \qquad \text{(Equation 2)}$$

For a given stimulus condition ($k$), $\theta_k$ and $r_k$ stands for the corresponding stimulus orientation (in radians) and the response of a unit. The unit was considered orientation selective for CV < 0.5 (an arbitrarily chosen threshold); otherwise, it was treated as orientation nonselective. Qualitatively similar results were observed for a large range of CV thresholds.

Before adding the hypothetical codes to the chosen layer (L4), the responses of all L4 units to an image were averaged and defined as the baseline modulation level (1 a.u.). The modulation levels of different codes were all normalized to this baseline level so that, at the same modulation level and in response to the same image, different codes displaced the population response vector by a fixed distance in the population space; the only difference among these codes was the direction of the displacement. Such a manipulation, which was restricted to L4 units in response to the object region only, took a modulation level from −0.5 to 4.5 in 0.1 steps. The influences of the three types of hypothetical codes (see Fig. 1E) on image classification were then tested at various modulation levels (Fig. 6D).

The above hypothetical codes were implemented with different degrees of feature specificity (i.e., specificity to oriented cues examined in the current study). Available physiological evidence indicates that surface segmentation involves two types of neural codes dissociable in space and time: the code for boundary detection followed by the code for interior region filling-in (*16*). By introducing spatial restriction to the additive code implemented in the CNN, we demonstrated the advantage of the object-level code (a uniform increase in responses of all units covered by the object) over the boundary-specific code (additive effect restricted within 5 pixels on both sides of the object boundary) and the interior-specific code (additive effect confined to the interior region 5 pixels away from the boundary) at different modulation levels (Fig. 6E).

Considering the spatial heterogeneity of saliencies commonly seen in natural scene images (e.g., Fig. 3J), the additive code uniformly applied to the entire object region was compared with that exclusive of a number of (0 to 11) randomly-chosen boundary locations. The modulation at each location was multiplied by a 2D Gaussian weight (pedestal = 1; amplitude = −1; aspect ratio = 2:1, the SD along the major axis aligned with the boundary was 20% of the squared root of the object area). With these settings, the modulation at the Gaussian center was eliminated (see Fig. 6F insets for examples of local attenuations).

**Transformation of downstream representations**

To understand how the segmentation code in an early cortical area (e.g., V1) would influence downstream representations, we quantified—before and after inclusion of the additive code in L4—the representational similarity for the same group of CNN units in a downstream layer in response to the same set of images. Due to limited computational resources, we confined the subsequent analyses to 1,838 images assembled from the aforementioned ImageNet-S919 by randomly drawing two images from each of the 919 classes.

Take the last fully connected CNN layer as an example. Given a set of 1,838 images, we sampled all 4,096 units in this layer and performed PCA on the resulting 1,838 × 4,096 matrix. The first 500 PCs were kept, which explained ~99.9% of the total variance. A linear regression was built between the pair of 1,838 × 500 matrices obtained before and after the additive manipulation. The coefficient of determination ($R^2$) was regarded as the representational similarity, which quantified the fraction of shared variances between these two matrices.

We repeated the above representational similarity analysis for each layer downstream L4. The following three hypotheses were examined as to how the segmentation code added to L4 could impact the downstream representations.

We first examined the hypothesis of representation amplification, which predicts a simple scaling up of downstream representations after inclusion of the additive code in L4. We tested this hypothesis in each downstream layer by using the degraded images (30% degradation level) and measuring the representational similarity (see above) before and after inclusion of the additive code. If the segmentation serves to amplify downstream representations, comparable representational similarities should be observed at various modulation levels. This is inconsistent with the simulation results (fig. S10A).

We next examined the denoising hypothesis, which posits that the segmentation serves to reduce external noise and in effect to revert the impact of image degradation in downstream representations. We tested this hypothesis in each layer downstream L4 by computing the representational similarity between two conditions: the degraded images with the inclusion of the additive segmentation code in L4, and the original undegraded images without introducing the code. The hypothesis predicts an increase in representational similarity with increasing the segmentation modulation, inconsistent with the simulations in deep layers (fig. S10B).

Last, we examined the cropping hypothesis that the additive segmentation code in L4 allows downstream analyses to focus on the isolated object region. We tested this hypothesis in each downstream layer by measuring the representational similarity between two conditions: the intact images with the inclusion of the additive segmentation code in L4, and the cropped images without introducing the code. The prediction is an increase in representational similarity with the modulation level, as observed (Fig. 6G).

### Evolving the segmentation code

Is the additive code the optimal solution for image segmentation? We addressed this question by evolving segmentation solutions from the weight-fixed CNN. A modulation matrix ($256 \times 56 \times 56$) attached to the corresponding units in L4 was optimized to recapitulate the three observed effects of the additive code: an improved representational similarity to the cropped images in the last fully connected layer (see Fig. 6G; see also the cropping hypothesis in **Transformation of downstream representations**); an unchanged representational geometry for natural images in L4 before and after evolving the attached modulation matrix (akin to the consistent orientation manifold in V1 between the object-absent and object-centered conditions, Fig. 5E); and an improved classification performance (see Fig. 6C). A stochastic gradient descent optimizer with momentum (*65*) was employed to optimize the modulation matrix in 100 training epochs. During the training, only the modulation matrix was adjusted, leaving all the pretrained CNN weights unchanged. Note that the modulation matrix had much higher degrees of freedom compared with the pretrained L4 units for two reasons: no weight sharing forced across spaces, and no constraints from natural image statistics such as edge continuity.

After the training, statistics were conducted on the modulation matrix. For a given image, we first computed a map of modulations in L4 (Fig. 6, I to K, bottom right panels; see also fig. S10C) by averaging the modulation matrix along its first dimension, i.e., across the 256 convolutional filters (aka feature maps). A map of activations was also computed similarly using the baseline responses of L4 units without introducing the modulation matrix (Fig. 6, I to K, bottom left panels; see also fig. S10C). Pearson correlation between the two maps was calculated, which was further compared with the correlation between the modulation map and the ground truth segmentation mask (Fig. 6L).

To study the feature specificity (i.e., specificity to oriented cues) of the evolved segmentation code, we visualized the relationship between the activations and modulations of the same L4 units. The evolved modulations within the object region were first binned based on the percentiles of corresponding activations (Fig. 6M), and the Pearson correlation between the binned modulations and activations was then calculated. Next, the mean modulation was computed for each L4 unit by averaging over the object region and across images, and the relationship between the mean modulation and the orientation selectivity (Equation 2) was examined across individual units (Fig. 6N).

**References and Notes (46–65)**

51. D. P. A. Schulz, M. Sahani, M. Carandini, Five key factors determining pairwise correlations in visual cortex. *J. Neurophysiol.*, (2015).

52. S. Siegel, *Nonparametric statistics for the behavioral sciences*. (McGraw-Hill, New York, 1956), pp. xvii, 312.

53. D. L. Ringach, R. M. Shapley, M. J. Hawken, Orientation selectivity in macaque V1: diversity and laminar dependence. *J. Neurosci.* **22**, 5639-5651 (2002).

54. Student, Probable Error of a Correlation Coefficient. *Biometrika* **6**, 302-310 (1908).

55. D. J. Thomson, Spectrum Estimation and Harmonic-Analysis. *Proc. IEEE* **70**, 1055-1096 (1982).

56. W. S. Geisler, J. S. Perry, B. J. Super, D. P. Gallogly, Edge co-occurrence in natural images predicts contour grouping performance. *Vision Res.* **41**, 711-724 (2001).

57. I. Sobel, G. Feldman, *An Isotropic 3x3 Image Gradient Operator*. (1968).

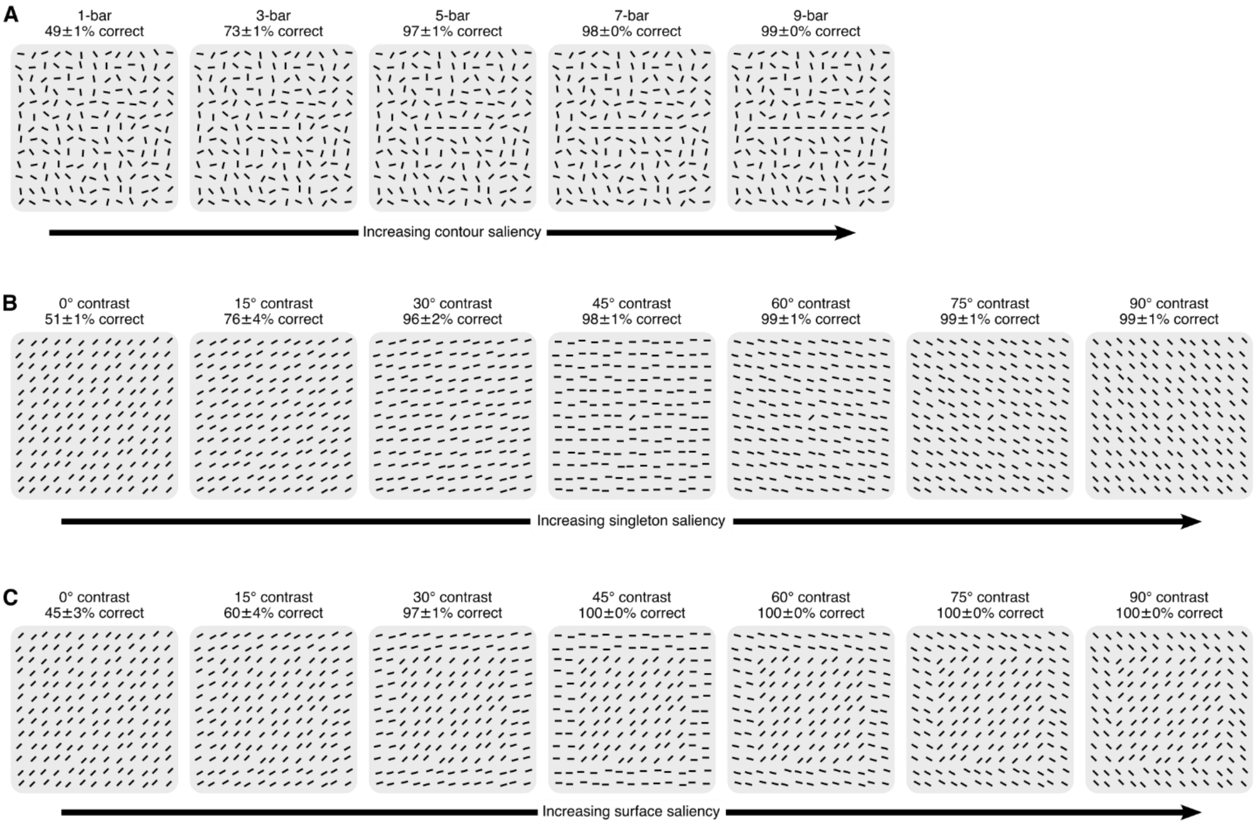


**fig. S1. Visual stimulus designs.**

**(A)** Examples of the contour stimuli. The stimuli were composed of short bars (0.25° × 0.05°) distributed in a hidden grid of 0.5° squares, each containing a bar with a random position jitter of up to 0.075° from the square center. A number of adjacent bars along a row of grids were collinearly aligned and evenly spaced, leaving the remaining bars at random orientations and jittered locations. The saliency of the contour object could be controlled by varying the number of collinear bars in the display. Captions on top of each panel indicate the saliency of the embedded object and the corresponding behavioral performance (mean ± SEM, trials from the two animals were pooled; 50% was the chance level, see fig. S2).

**(B)** Examples of the singleton stimuli. Among the short bars distributed in the hidden grid, the single bar defining the object and those belonging to the background were respectively arranged at two orientations. The saliency of the object could be controlled by adjusting the angular difference (i.e., orientation contrast) between the object and background bars.

**(C)** Examples of the surface stimuli. The design was the same as the singleton stimuli shown in B except that the singleton was replaced by 9 × 9 iso-oriented bars.

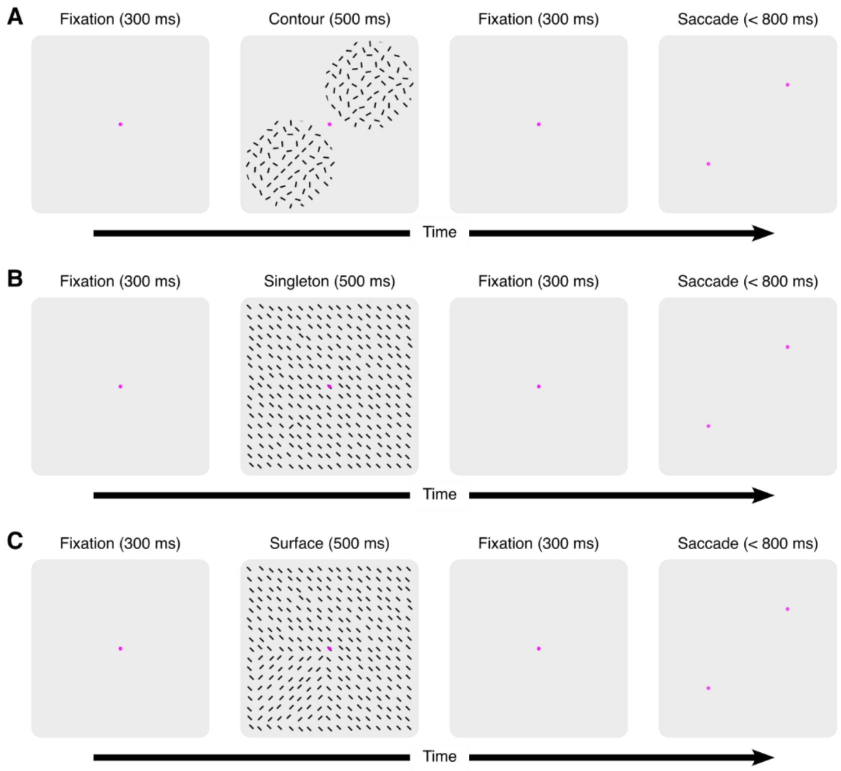


**fig. S2. Behavioral paradigms.**

Schematic of a representative trial in the contour (A), singleton (B), and surface (C) segmentation tasks. The trial began when the monkey fixated the central point. After 300 ms of fixation, the stimulus appeared for 500 ms, in which an object (contour, singleton, or surface) was embedded in the background randomly at one of two locations symmetric around the fixation point. Subsequently the stimulus disappeared, and two dots were presented at the two locations. The monkey was rewarded for making a saccade within 800 ms to the dot corresponding to the location where the object had been shown.

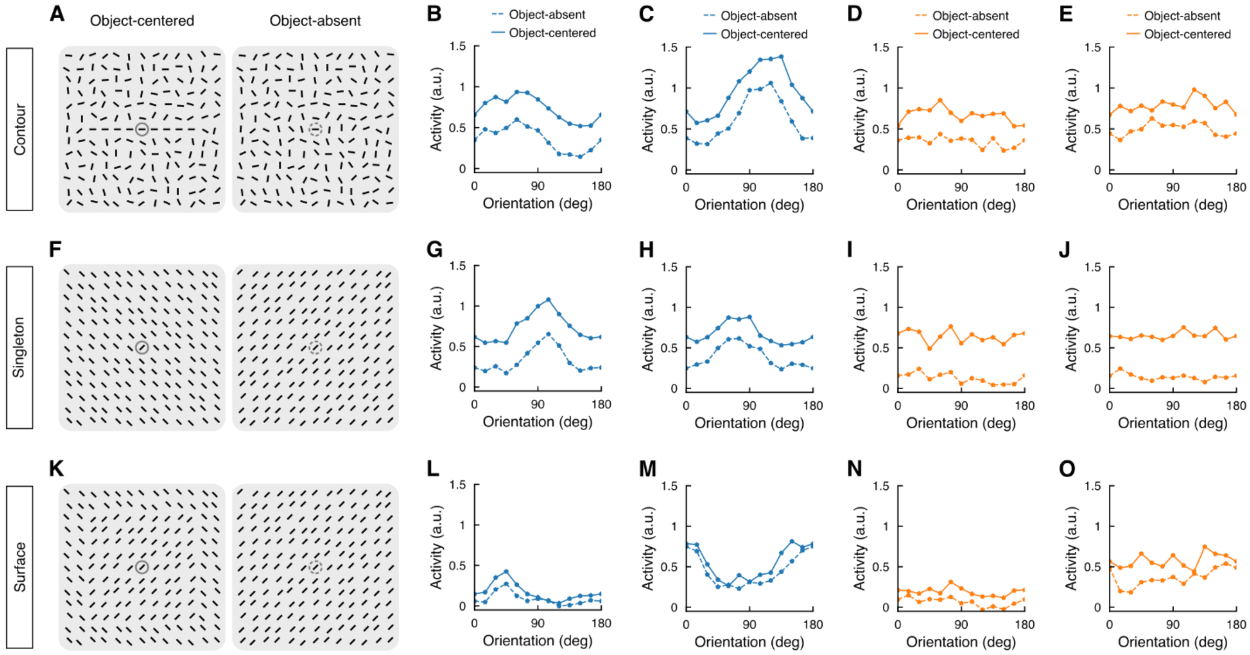


**fig. S3. Elevation of the orientation tuning curves of sample V1 sites during segmentation.**

**(A)** Contour stimuli in the object-centered (left) and object-absent (right) conditions, with the RF indicated by the gray circle. The orientation tuning curves of a V1 site were measured by rotating the stimulus pattern from 0° to 180° in 15° steps under these two conditions.

**(B** to **E)** Normalized tuning curves of two orientation-selective (B and C) and two nonselective (D and E) V1 sites in the object-absent (dashed line) and object-centered (solid line) conditions. The contour centered on the RF resulted in an additive increase in the tuning curve regardless of neuronal orientation selectivities.

**(F** to **O)** Similar to A to E, for the singleton (F to J) and surface (K to O) stimuli. The orientation tuning curves from different examples of V1 sites were measured by simultaneously rotating all of the bars by the same amount around their respective centers, keeping the object outline and saliency unchanged.

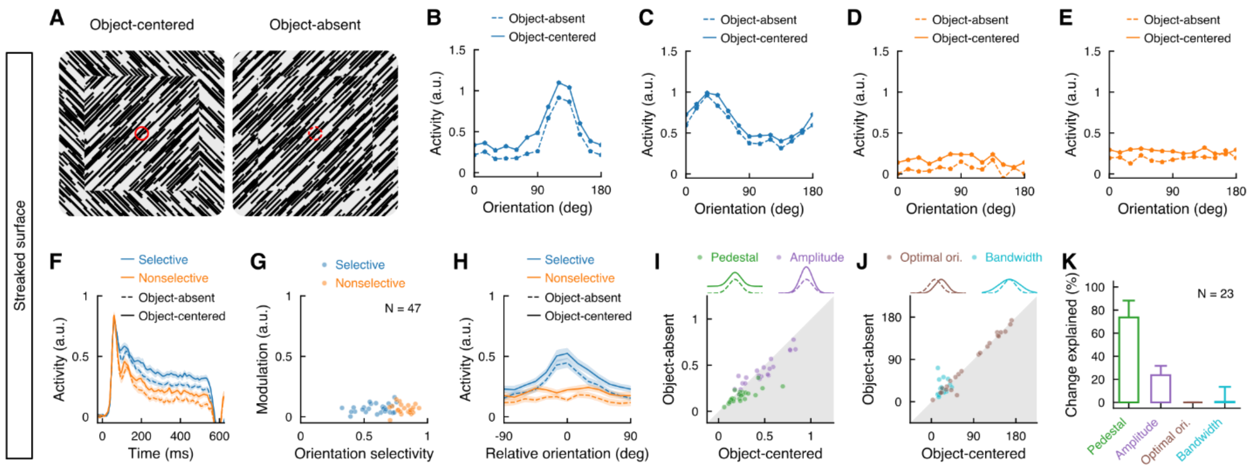

**fig. S4. Additive code in V1 for the streaked surface.**

**(A** to **E)** Sample stimuli of the streaked surface (A), and the tuning curves of two orientation-selective (B and C) and two nonselective (D and E) V1 sites.

**(F)** Population PSTHs in response to the streaked surface in the object-absent (dashed) and object-centered (solid) conditions, averaged across all the orientations and plotted separately for orientation-selective (blue) and nonselective (orange) V1 sites. Color shades indicate standard errors of the mean (SEM); time 0, the stimulus onset.

**(G)** Comparison of the segmentation signal strength (i.e., modulation) and orientation selectivity across individual V1 sites. No significant correlation was observed ($\rho = 0.07$, $P = 0.63$).

**(H)** Population-averaged orientation tuning curves in the object-absent and object-centered conditions for orientation-selective and nonselective V1 sites, respectively.

**(I** and **J)** Comparisons of the tuning parameters (from Gaussian fit, Equation 1) for orientation-selective V1 sites in the object-absent and object-centered conditions. Introducing an object to the RF location increased the tuning pedestal ($P < 10^{-4}$) and amplitude ($P = 0.02$), but did not alter the optimal orientation ($P = 0.26$) and bandwidth ($P = 0.60$) of the tuning curves.

**(K)** Contribution of individual tuning parameters to tuning curve changes between the object-absent and object-centered conditions. The additive code, i.e., an elevated tuning pedestal, explained most of the orientation tuning changes (74%).

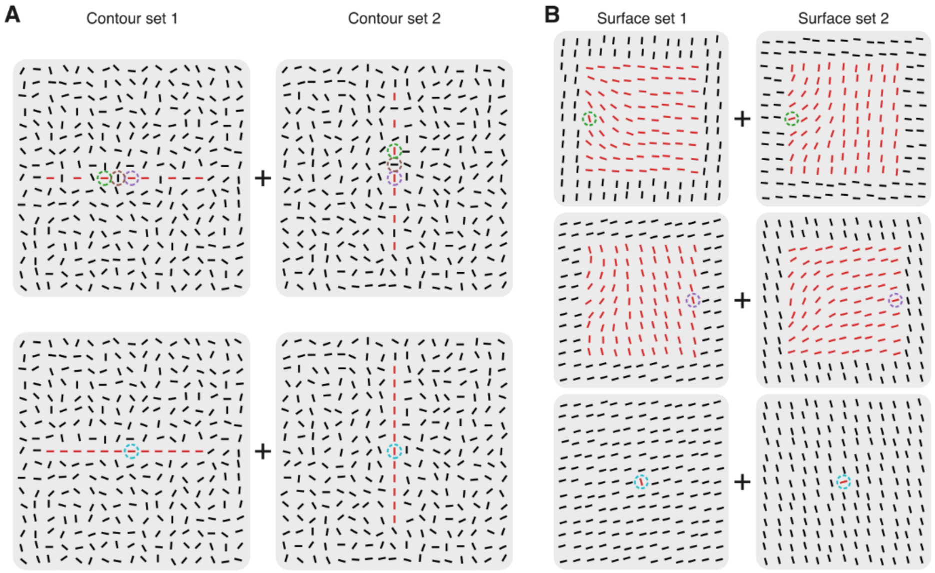


**fig. S5. Complementary stimulus sets used in the object-level saliency experiments.**

**(A)** Two complementary sets of chimera contours (top, left + right) used to balance the responses of a V1 site. The contour path was aligned with, or orthogonal to, the optimal orientation the recorded site. The RF could be centered on any of the component bars in the middle triplet (color circles indicate 3 possible RF locations). As a control, we also tested the condition that all component bars of the chimera contour were collinearly aligned (bottom).

**(B)** Two complementary sets of chimera surface used to balance the responses of a V1 site. The difference between each pair of stimuli (left vs. right) was a rotation of all the bars by 90° around their respective centers so that the bar in the RF (small circle) could be optimally (left) or orthogonally (right) oriented. The RF was placed respectively at two corresponding boundary locations where the orientation contrast between the bar in the RF and the iso-oriented background bars was 20° (top) or 90° (middle). The singleton stimulus (bottom) was used as a control.

The simple objects (contours, surfaces and singletons) are highlighted in red simply for illustrative purposes.

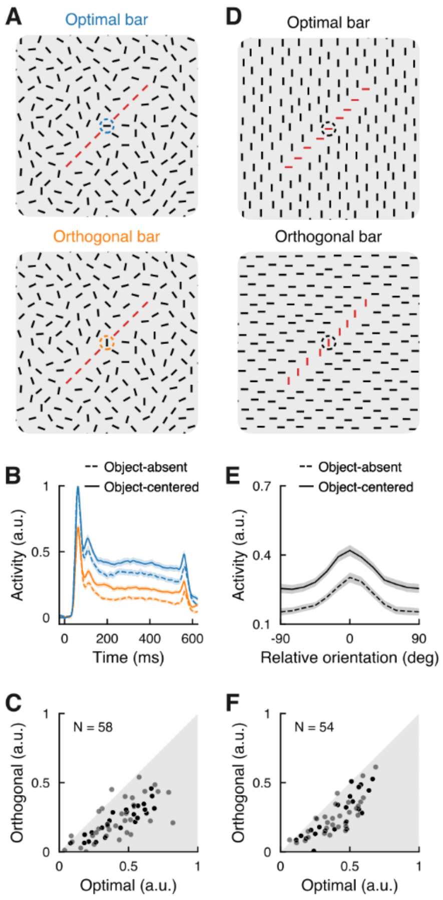


**fig. S6. Neuronal selectivity for local stimulus orientations in the absence or presence of a global object.**

**(A)** Examples of the contour stimuli, in which the local orientation of the bar in the RF was varied while the contour orientation remained unchanged (upper vs. lower panel).

**(B)** Population PSTHs in response to the contour stimuli when the bar in the RF was optimally (blue) and orthogonally (orange) oriented in the object-absent (dashed) and object-centered (solid) conditions, respectively. The same V1sites in Fig. 4, A to C, were included in this analysis.

**(C)** Comparison of the mean responses of individual V1 sites to the optimally and orthogonally oriented bar in the absence of the contour (i.e., object-absent condition in B). Light and dark points represent data from the two animals, respectively.

**(D)** Examples of the slim surface stimuli, in which the local orientation of the surface component bars was varied while the surface orientation and the surface-background orientation contrast remained unchanged (upper vs. lower panel).

**(E)** Population averaged tuning curves for the local bar orientations in the object-absent (dashed) and object-centered (solid) conditions for the slim surface stimuli. The same V1 sites in Fig. 4, D to F, were included in this analysis.

**(F)** Comparison of the mean responses of individual V1 sites to the optimally and orthogonally oriented bar in the absence of the slim surface (i.e., object-absent condition in E).

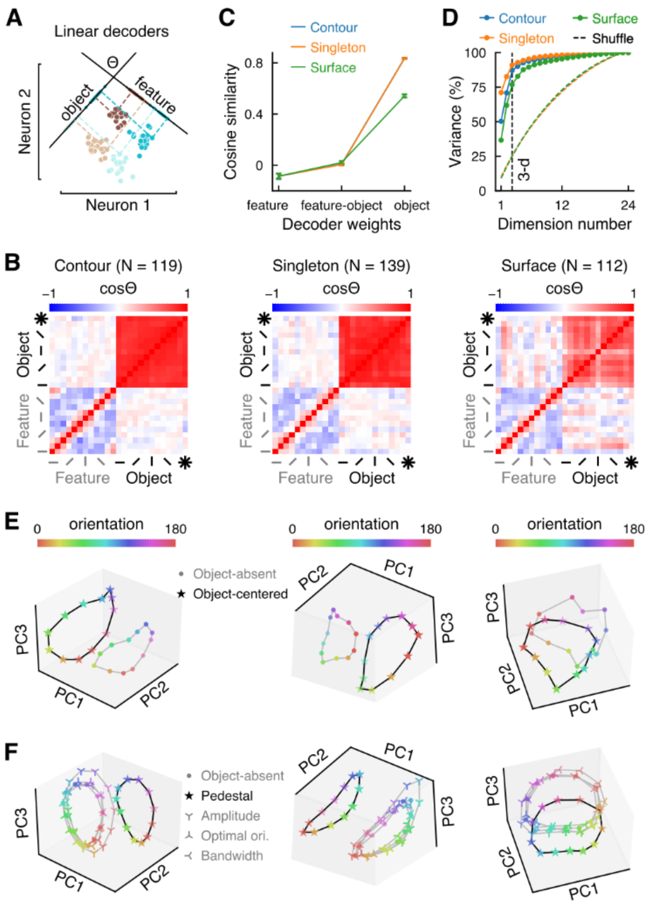


**fig. S7. Neural manifold visualization based on all recorded V1 sites.**

Companion visualizations of Fig. 5, computed from the entire neuronal population by pooling both orientation-selective and nonselective V1 sites. All the individual panels correspond to those in Fig. 5, showing a robust representational geometry in different segmentation scenarios (contour, singleton, and surface) across the entire population of recorded V1 sites (this figure) or a subset of them (i.e., orientation-selective sites in Fig. 5).

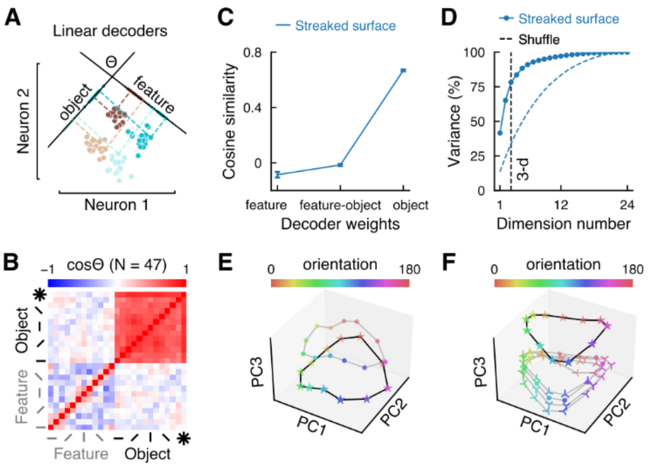


**fig. S8. Neural manifold visualization for the streaked surface.**

Companion visualizations of fig. S7 and Fig. 5, computed from the streaked surface dataset from the orientation independency experiments (fig. S4). Orthogonality between the feature and object decoders was also observed in this dataset (B and C), which was further consolidated by the orthogonal displacement of the ring-shaped manifold for stimulus orientations (parallel to the plane defined by PC1 and PC2) along the third dimension of PC3 (E). The orthogonal displacement was well explained by an increase in the tuning pedestal (F, star; parameter $b$ in Equation 1) of the orientation tuning curve, i.e., an additive code in the population space. This representational geometry for the streaked surface is consistent with that for the surface formed by 9 × 9 bars (see Fig. 5, E and F, right panels).

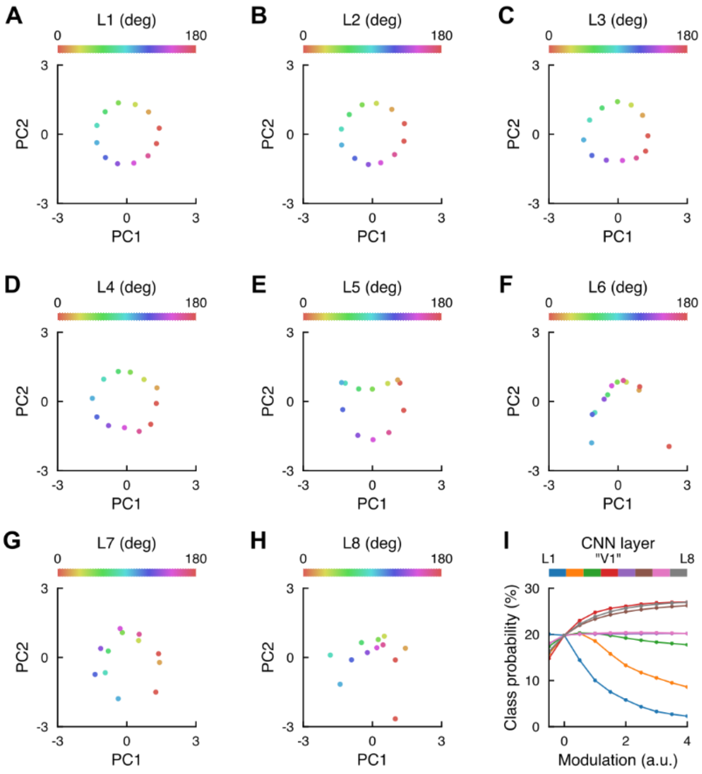


**fig. S9. CNN layer selection.**

**(A** to **H)** Visualization of the orientation manifold in different convolutional layers (L1 to L8) in the CNN model. PCA was applied to the units' responses of a given convolutional layer to gratings of different orientations (color coded). The orientation manifold for the given layer was visualized based on its embeddings along the first two PCs. According to the degree of similarity to macaque V1 (in comparison with the ring-shaped orientation manifolds shown in Fig. 5E), L1 to L4 were considered as the candidates for the V1 counterpart in the CNN.

**(I)** Model performance as a function of the segmentation modulation level. Implementation of the additive code in different convolutional layers (denoted by colors) showed different impacts: deterioration for L1-3 but improvement for L4, L6 and L8, with the largest improvement observed in L4. Together with the other two criteria (see **Layer selection**), L4 was chosen as the V1 counterpart in the CNN (labeled "V1" here). All subsequent manipulations in the computational modeling targeted this layer.

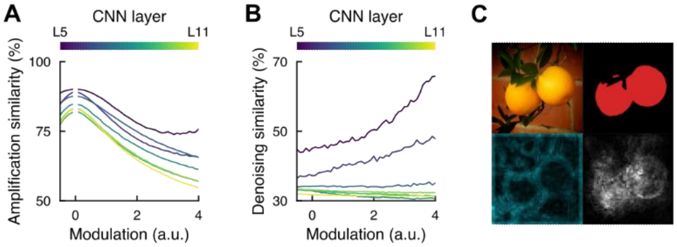


**fig. S10. More characteristics of the segmentation modeling.**

**(A)** Simulation test for the representation amplification hypothesis, which predicts a simple scaling up of downstream representations after inclusion of the additive code in L4. Representational similarity before and after the manipulation of L4 was quantified across modulation levels for each downstream layer (color coded). If the segmentation serves to amplify downstream representations, comparable representational similarities should be observed at various modulation levels. This is inconsistent with the simulation results.

(B) Simulation test for the denoising hypothesis, which posits that the segmentation serves to reduce external noise and in effect to revert the impact of image degradation in downstream representations. For each layer downstream L4, we computed the representational similarity between two conditions: the degraded images with the inclusion of the additive segmentation code in L4, and the original undegraded images without introducing the code. The hypothesis predicts an increase in the representational similarity with increasing the segmentation modulation, inconsistent with the simulations in deep layers.

**(C)** An additional example of evolved modulations (similar to Fig. 6, I to K). The evolved modulation map in L4 (bottom right) is more similar to the ground truth object mask (top right) than the map of unit activations (bottom left).
